# Angptl5 restricts primitive hematopoiesis by modulating retinoic acid signaling in zebrafish

**DOI:** 10.1101/2025.08.18.670818

**Authors:** Jing Mo, Ding-Hao Zhuo, Ying Huang, Tao Cheng, Yang Dong, Yan-Yi Xing, Yun-Fei Li, Zi-Xin Jin, Xiang Liu, Guo-Qin Zhao, Hai-Rong Pu, Yu-Meng Liu, Zhi-Xu He, Li-Ping Shu, Peng-Fei Xu

**Affiliations:** Department of Immunology, School of Basic Medicine, Guizhou Medical University, Guiyang, Guizhou, China, 561113; Center for Genetic Medicine, the Fourth Affiliated Hospital of School of Medicine, and International School of Medicine, Zhejiang University, Yiwu, China, 322000; Institute of Genetics, Zhejiang University School of Medicine, Hangzhou, Zhejiang, China, 310063; Women’s Hospital, Zhejiang University School of Medicine, Hangzhou, Zhejiang, China, 310006; National & Guizhou Joint Engineering Laboratory for Cell Engineering and Biomedicine Technique, and Guizhou Province Key Laboratory for Regenerative Medicine, Guizhou Medical University, Guiyang, Guizhou, China, 561113

**Keywords:** Angptl5, primitive hematopoiesis, Integrin α6l/β5, retinoic acid (RA), Dhrs9

## Abstract

Homeostasis is essential for hematopoiesis, and its dysregulation can lead to severe pathological conditions. Retinoic acid (RA) is a key regulator that exerts concentration-dependent effects on both embryonic and adult hematopoiesis. However, the mechanisms that modulate RA signaling in hematopoietic processes remain poorly understood. Using zebrafish as a model, we identified angiopoietin-like protein 5 (Angptl5) as a critical regulator of hematopoietic homeostasis. Loss of Angptl5 function resulted in myeloid hyperplasia in the anterior lateral plate mesoderm (ALPM) and anterior expansion of erythroid progenitors in the posterior lateral plate mesoderm (PLPM)— phenotypes consistent with attenuated RA signaling. Molecular analyses confirmed impaired RA signaling in *angptl5^Δ10/Δ10^* mutants, and exogenous RA supplementation fully rescued the hematopoietic defects. Mechanistically, we found that Angptl5 transcriptionally activates retinol dehydrogenase *dhrs9* through its interaction with Integrin α6lβ5. Our findings establish Angptl5 as a novel and essential regulator of embryonic hematopoiesis and reveal a previously unrecognized mechanism controlling hematopoietic homeostasis. These insights position Angptl5 as a potential therapeutic target for hematological disorders.

## Introduction

Primitive hematopoiesis in vertebrates, such as zebrafish, is spatially organized along the anterior-posterior (A-P) embryonic axis, with distinct progenitor populations emerging from the anterior lateral plate mesoderm (ALPM) and posterior lateral plate mesoderm (PLPM) (1–3). The transcription factor *tal1* serves as a critical marker for the onset of the hematopoietic program, with expression initiating in the ALPM and PLPM at the 2-somite stage in zebrafish (4). In the ALPM, subsets of *tal1*-expressing cells acquire a myeloid fate, marked by the expression of *spi1b* (a myeloid-specific transcription factor) (5). In contrast, the PLPM exhibits bipotent erythro-myeloid potential, as evidenced by the co-expression of *spi1b* and *gata1a* (an erythroid-specific transcription factor) (6–8).

Retinoic acid (RA), a key morphogen involved in patterning the A-P axis and regulating tissue specification during embryonic development, has been shown to suppress *spi1b* expression in the ALPM (9). Interestingly, RA inhibition causes an anterior shift of *gata1a*-positive cells (10), suggesting differential RA sensitivity between compartments (11). These observations imply that RA function as a molecular “rheostat” in regulating myeloid and erythroid fates. In addition, RA has been successfully used to treat specific types of leukemia (12). However, the mechanism modulating RA in orchestrating primitive hematopoiesis remains unclear.

Angiopoietin-like proteins (ANGPTLs) are a family of secreted glycoproteins involved in a variety of developmental and disease processes, including angiogenesis, hematopoietic stem cells maintenance and cancer progression (13, 14). For example, ANGPTL1 suppresses the integrin signaling to inhibit angiogenesis and metastasis in hepatocellular carcinoma (15), while ANGPTL4 is essential for common myeloid progenitor differentiation and inflammatory responses (16, 17). ANGPTL5 has been shown to enhances *ex vivo* expansion of human cord blood hematopoietic stem cells (HSCs) (18) and interact with inhibitory receptors to support leukemia progression (19), however, its function during embryonic development remains unexplored.

In this study, we used zebrafish as a model to investigate the function of Angplt5 in primitive hematopoiesis. We found that *angptl5* knockout results in myeloid hyperplasia in the ALPM and anterior expansion of erythroid progenitors in the PLPM-phenotypes reminiscent of RA deficiency. Molecular analysis revealed downregulation of RA signaling in angptl5 mutants, and RA treatment was able to rescue the hematopoietic defects. Mechanistically, we found that Angplt5 transcriptionally activates the retinol dehydrogenase *dhrs9*, a key enzyme in the RA synthesis pathway, through its interaction with integrin α6lβ5 and subsequent activation of the ERK signaling pathway. Collectively, our findings identify Angptl5 as a new regulator of primitive hematopoiesis that function upstream of RA signaling.

## Results

### *angptl5* is expressed in the primitive hematopoietic tissue during zebrafish embryonic development

Angiopoietin-like proteins are secreted glycoproteins conserved across vertebrates (Supplemental Figure 1A-B). Members of this family feature two highly conserved domains: an N-terminal coiled-coil domain (CCD) and a C-terminal fibrinogen-like domain (FLD) (Supplemental Figure 1C). To investigate their potential roles in development, we first analyzed the spatial and temporal expression patterns of angiopoietin-like genes during zebrafish embryogenesis using whole-mount *in situ* hybridization (WISH). Interestingly, different angiopoietin-like genes exhibited distinct tissue specificity. For example, *angptl2b* was expressed in the posterior spinal cord, *angptl6* in the notochord, and *angptl7* in the somites (Supplemental Figure 2A). Among these, *angptl5* was expressed in the mesendoderm during gastrulation and specifically enriched in hematopoietic tissues at 24 hours post-fertilization (hpf), including the rostral blood island (RBI) and posterior intermediate cell mass (ICM) (Figure 1A, Supplemental Figure 2B-C). To further confirm its expression, we generated a transgenic line *Tg*(*angptl5:EGFP*) by cloning the upstream 867 bp region of *angptl5* gene and EGFP coding sequece, which revealed expression of *angptl5* in the ALPM and partial co-localization with *spi1b*-positive cells (Figure 1B, Supplemental Figure 2D). Additionally, analysis of a single-cell RNA sequencing (scRNA-seq) dataset showed that *angptl5* is expressed in a subset of *spi1b*-positive cells at both 18 hpf and 24 hpf (Figure 1C-F) (20). Together, these findings suggest that *angptl5* may play a role in primitive hematopoiesis in zebrafish.

**Figure 1.**
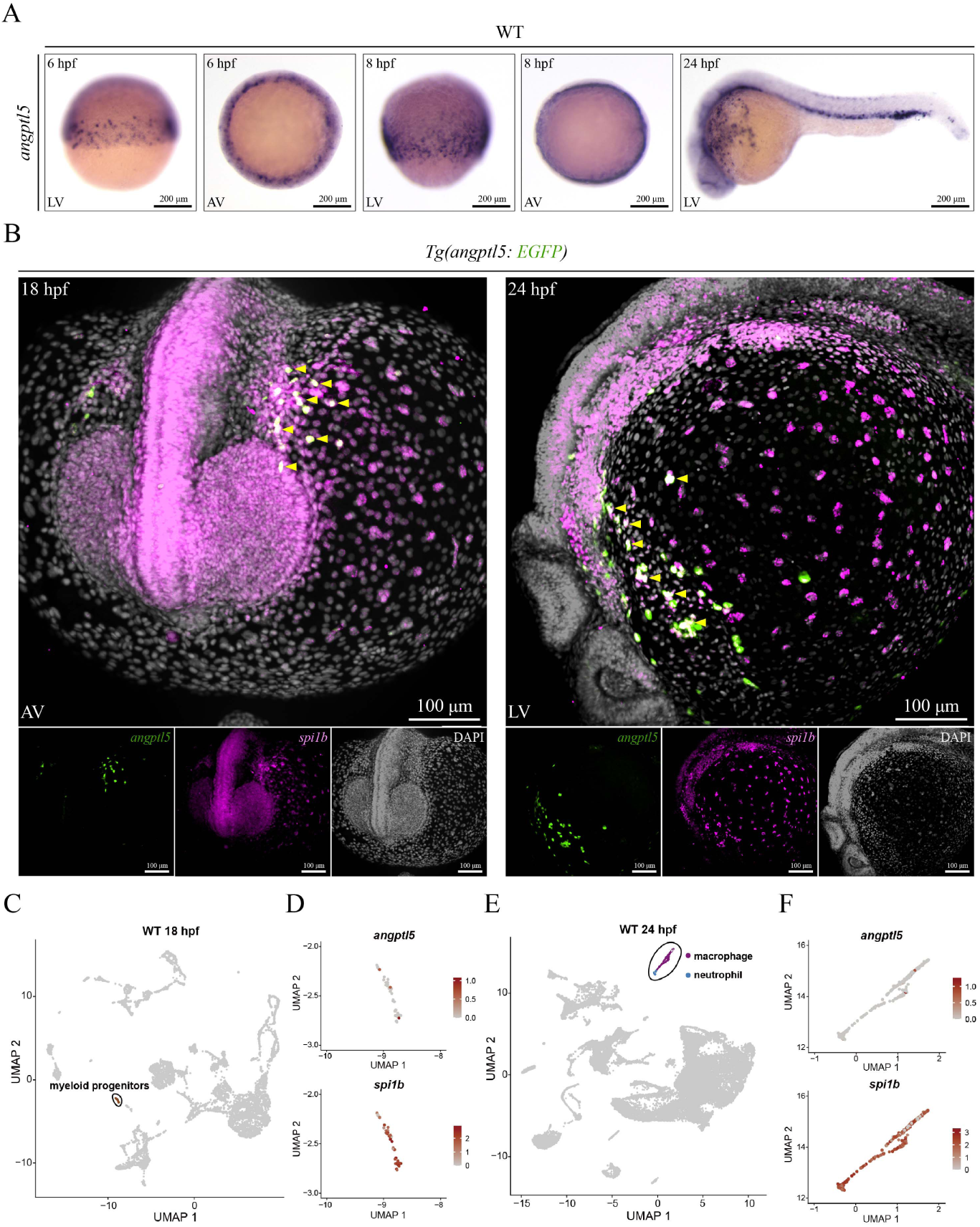
Expression pattern of *angptl5* during zebrafish embryonic development. (A) Whole-mount in situ hybridization (WISH) of *angptl5* in zebrafish embryos at the indicated stages. (B) Projection of images from confocal stacks to show co-localization of myeloid progenitor marker *spi1b* mRNA and EGFP protein in *Tg(angtl5:EGFP)*. Each imaging was performed for at least three independent replicates. (C-F) Co-expression of *angptl5* and *spi1b* in single-cell analysis. UMAP analysis of 18 hpf (C) and 24 hpf (E) WT zebrafish embryos from the published dataset(20), with myeloid lineage cells highlighted. Feature plots showing the expression of *angptl5* and *spi1b* in myeloid progenitors (D) and in myeloid lineages (F). hpf, hours post-fertilization; LV, lateral view; AV, animal view.

### *angptl5* mutation leads to hyperplasia of myeloid and erythroid progenitors

To investigate the function of Angptl5 in hematopoiesis, we generated two *angptl5* mutant lines using CRISPR/Cas9: *angptl5Δ10* (10 bp deletion) and *angptl5Δ5* (5 bp deletion) (Supplemental Figure 3A). Both mutations induced frameshifts producing premature stop codons, resulting in truncated proteins that either lacked the fibrinogen-like domain (FLD; Angptl5Δ10) or both the coiled-coil domain (CCD) and FLD (Angptl5Δ5).

While expression of hemangioblasts marker *tal1* appeared normal, *angptl5Δ10* mutants displayed significant expansion of *spi1b*+ myeloid progenitors in the ALPM (Figure 2A-B), along with increased neutrophils (*lyz*+, *mpx*+) and macrophages (*mpeg1.1*+) (Supplemental Figure 3B). This phenotype was consistently observed in *angptl5Δ5* mutants (Supplemental Figure 3C-D), suggests that loss of angptl5 function induces defects in primitive myelopoiesis. Based on these findings, the *angptl5Δ10* mutant was selected for further functional analysis. Markedly, ectopic *angptl5* mRNA expression effectively suppressed the aberrant expansion of *spi1b+* myeloid progenitor in *angptl5^Δ10/Δ10^* embryos (Figure 2C). We then checked the expression pattern of *gata1a*+ erythroid progenitors. Interestingly, HCR co-staining with the somite marker *myod1* revealed anterior expansion of *gata1a* (Figure 2D), a phenotype resembling attenuated RA signaling (10). And quantitative real-time PCR (qPCR) also confirmed upregulation of *spi1b* and *gata1a* in *angptl5^Δ10/Δ10^* embryos, while *tal1* and the angioblast marker *etsrp* remained unchanged (Supplemental Figure 3E).

**Figure 2.**
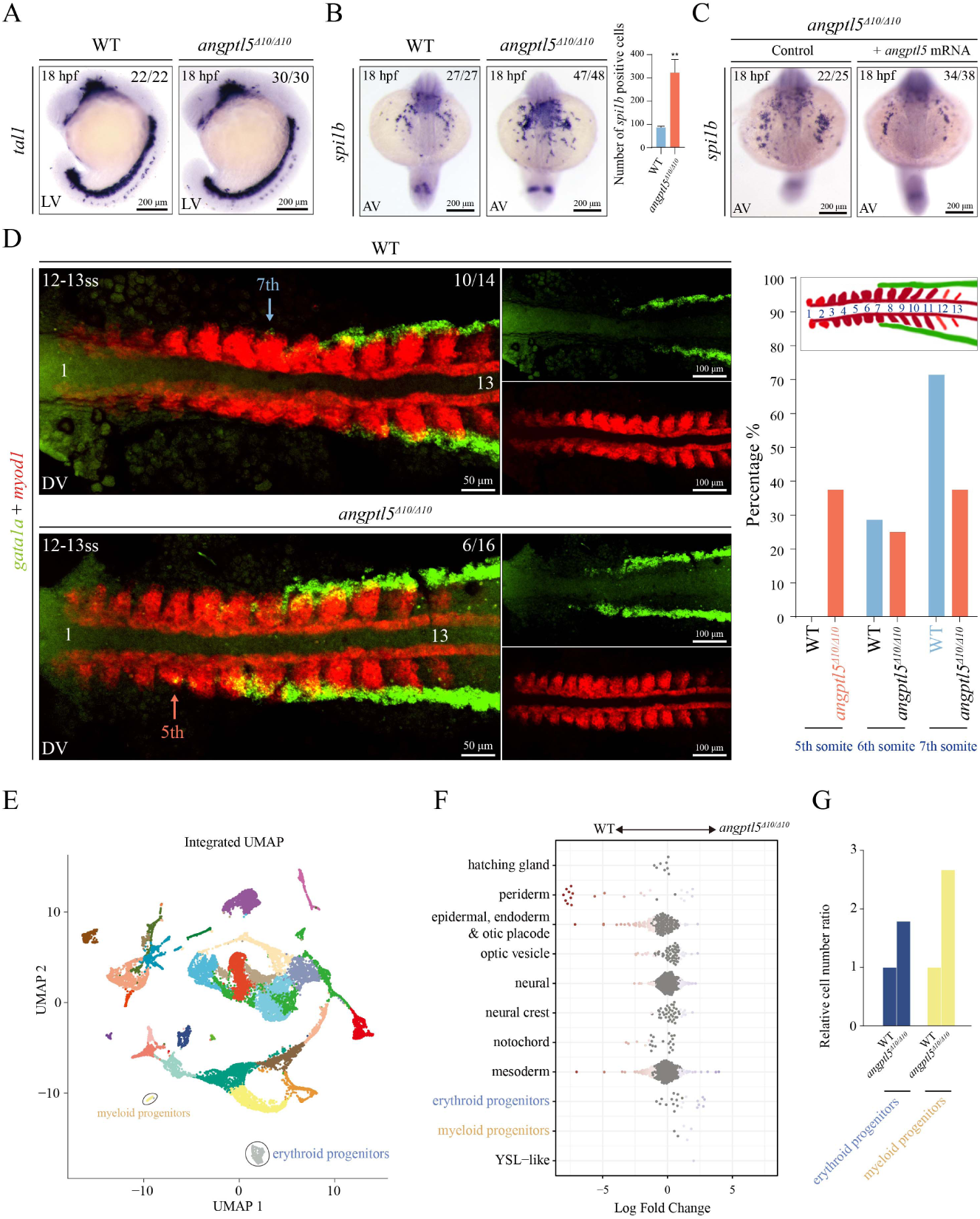
Angptl5 deficiency impairs primitive hematopoiesis. (A-B) WISH of *tal1* (A) and *spi1b* (B) in WT and *angptl5^Δ10/Δ10^* embryos. Data presented as the mean ± SD. Statistical significance: **P < 0.01. (C) WISH of *spi1b* in *angptl5^Δ10/Δ10^* embryos injected with *angptl5* mRNA at the 1-cell stage. Uninjected embryos were used as control. (D) Projection of images from confocal stacks to show *gata1a* and *myod1* expression in flat-mounted embryos in WT and *angptl5^Δ10/Δ10^*embryos as assessed using HCR. Statistics are shown on the right. (E) Integrative UMAP analysis of 16 hpf WT and *angptl5^Δ10/Δ10^* embryos. The WT dataset is from the published work(21). Each cell is coloured according to cell type annotations. (F) Bees warm plot showing the differential abundance by cell types between WT and *angptl5^Δ10/Δ10^*embryos. (G) Bar plot showing the relative cell number ratios of each hematopoietic subtype to the total cell count in *angptl5^Δ10/Δ10^* embryos compared to WT embryos. LV, lateral view; AV, anterior view.

To further characterize the hematopoietic phenotype caused by Angptl5 deficiency, we performed single-cell RNA sequencing (scRNA-seq) on 16 hpf *angptl5^Δ10/Δ10^* mutant embryos and integrated these data with published WT scRNA-seq datasets (21). While UMAP visualization revealed similar overall cell type distributions between mutant and WT embryos (Figure 2E, Supplemental Figure 4A-B), differential abundance analysis identified significant expansion of hematopoietic populations in mutants, including erythroid progenitors and myeloid progenitors (Figure 2F-G).

Angiopoietin-like proteins have been previously reported to play roles in angiogenesis (22, 23); however, using either WISH or transgenic reporter lines, we did not observe obvious changes in the vascular progenitor marker *etsrp* (24) or the endothelial marker *kdrl* (25) between WT and *angptl5^Δ10/Δ10^*embryos at 3 dpf and 3.5 dpf (Supplemental Figure 5A-C).

Taken together, the main phenotypes observed in *angptl5* mutants are myeloid hyperplasia in the ALPM and anterior expansion of erythroid progenitors in the PLPM, while the vasculogenesis remains largely unchanged.

### *angptl5* mutation leads to attenuated retinoic acid (RA) signaling

The elevated *spi1b* and anterior expansion of *gata1a* seen in *angptl5* mutants resemble the phenotype observed upon RA signaling inhibition in zebrafish (10) and mirror vitamin A deficiency-induced promotion of hematopoietic stem cell differentiation in mice (26). To determine whether RA signaling is affected in *angptl5* mutants, we firstly examined the expression of RA direct target *cyp26a1* (27), as well as retinol dehydrogenase *dhrs9* and aldehyde dehydrogenase *aldh1a2*, two key enzymes required for RA biosynthesis (Supplemental Figure 6A). Interestingly, WISH and qPCR analyses revealed significantly reduced expression of *cyp26a1* at 8 hpf and 18hpf, *aldh1a2* at 8hpf, and *dhrs9* at 8 hpf and 18hpf in *angptl5^Δ10/Δ10^*embryos (Figure 3A-C). Supporting these findings, *angptl5^Δ10/Δ10^*; *Tg*(*RARE*:EGFP) embryos exhibited markedly reduced EGFP fluorescence compared to *Tg(RARE:*EGFP) embryos, providing further evidence of attenuated RA signaling in *angptl5* mutants (Figure 3D, Supplemental Figure 6B).

**Figure 3.**
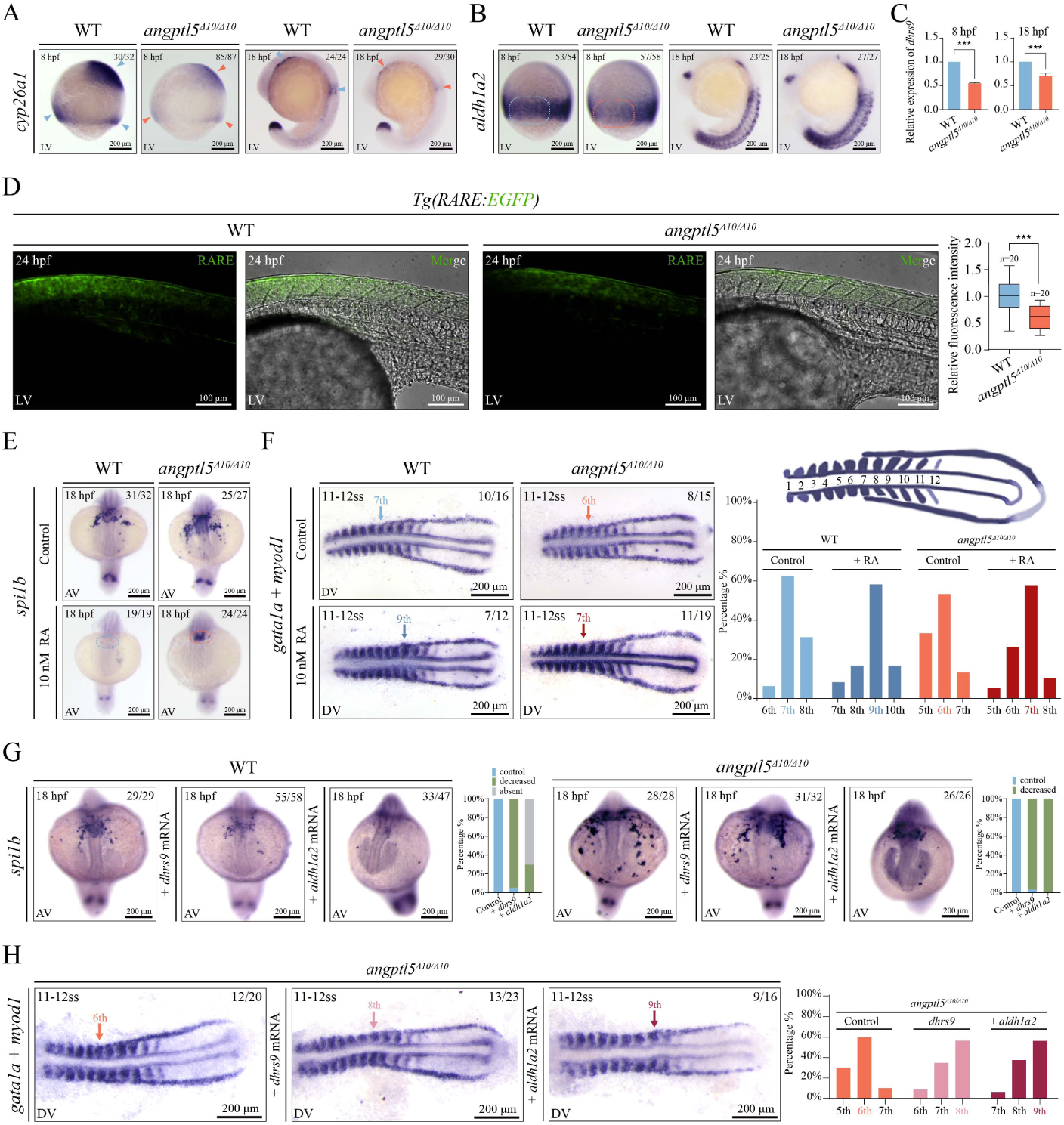
***angptl5* loss-of-function attenuates retinoic acid signaling**. (A-B) WISH of *cyp26a1* (A) and retinal dehydrogenase *aldh1a2* (B) in WT and *angptl5^Δ10/Δ10^* embryos. (C) Quantitative real-time PCR (qPCR) shows the retinol dehydrogenase *dhrs9* mRNA expression in WT and *angptl5^Δ10/Δ10^*embryos. Data presented as the mean ± SD. Three independent biological replicates were used. Statistical significance: ***P < 0.001. (D) Retinoic acid (RA) signaling in WT and *angptl5^Δ10/Δ10^* embryos shown by *RARE:EGFP* reporter. Statistics are shown on the right. Data presented as the mean ± SD. Statistical significance: ***P < 0.001. (E) WISH of *spi1b* in WT and *angptl5^Δ10/Δ10^* embryos treated with RA. Untreated embryos were used as control. (F) WISH of *gata1a* and *myod1* in flat-mounted WT and *angptl5^Δ10/Δ10^* embryos treated with RA from the shield stage to the 11-12 somite stage (ss). Statistics are shown on the right. (G) WISH of *spi1b* in WT and *angptl5^Δ10/Δ10^* embryos injected with *dhrs9* or *aldh1a2* mRNA at the 1-cell stage. Uninjected embryos were used as control. Statistics are shown on the right of the representative photos. (H) WISH of *gata1a* and *myod1* in flat-mounted WT and *angptl5^Δ10/Δ10^* embryos injected with *dhrs9* or *aldh1a2* mRNA at the 1-cell stage. Statistics are shown on the right. LV, lateral view; AV, anterior view; DV, dorsal view.

To investigate the hierarchy relation between Angptl5 and RA signaling in hematopoiesis regulation, we treated WT and *angptl5^Δ10/Δ10^*embryos with RA inhibitor (DEAB, inhibitor of *aldh1a2*) or RA at varying concentrations starting from shield stage. Consistent with previous studies, RA exhibited dose-dependent teratogenic effects, with concentrations ≥100nM causing severe morphological abnormalities in WT embryos (Supplementary Figure 6C) at 18 hpf. Pharmacological inhibition of RA markedly suppressed *cyp26a1* expression, whereas exogenous RA administration robustly upregulated this RA target gene in a dose-dependent manner (Supplementary Figure 6D). While RA treatment suppressed *spi1b* expression in all embryos, *angptl5* mutants demonstrated significantly higher resistance to RA treatment, as evidenced by persistent *spi1b* and *gata1a* expression compared to WT controls (Figure 3E-F, Supplementary Figure 7A). Consistently, treatment with the RA receptor antagonist AGN 193109 (AGN) or DEAB led to a significant expansion of *spi1b*^+^ and *gata1a*^+^ cells in both WT and *angptl5^Δ10/Δ10^*embryos (Supplementary Figure 7B-C).

Above results suggest that RA activation could compensate for Angptl5 deficiency. To further confirm the effect of restoring RA signaling on primitive hematopoiesis in *angptl5* mutants, we performed rescue experiments by injecting *dhrs9* or *aldh1a2* mRNA at the 1-cell stage in both WT and *angptl5^Δ10/Δ10^* embryos. Overexpression of both *dhrs9* and *aldh1a2* enhance the *cyp26a1* expression, indicating the elevated RA signaling (Supplementary Figure 7D). Consistently, the expanded expression of *spi1b*^+^ myeloid and *gata1a^+^*erythroid progenitors in *angptl5* mutants was rescued (Figure 3G-H). Together, these results indicate that Angptl5 function as an upstream regulator of RA-mediated hematopoietic patterning.

### Angptl5 physically interact with integrin α6lβ5

As a secreted protein, Angptl5 requires receptor binding to exert its biological functions. To comprehensively identify receptors through which Angptl5 regulates RA signaling, we employed TurboID-mediated proximity labeling combined with immunoprecipitation-mass spectrometry (IP-MS) (28) in zebrafish. We engineered a construct expressing *angptl5* mRNA fused with a C-terminal Flag-tagged TurboID. Secretion of engineered proteins were detected in HEK293T conditioned medium and the expression of tagged *angptl5* and biotinylated proteins were detected in zebrafish embryos (Supplemental Figure 8A-B). The engineered *angptl5* mRNA exhibited similar rescue efficiencies to untagged controls, indicating that the fusion tags do not compromise Angptl5 protein functionality (Supplemental Figure 8C). We then injected engineered WT and mutated *angptl5* mRNA into 1-cell stage embryos, and incubate those embryos with biotin for 8 hours before collecting them for IP-MS, respectively (Figure 4A).

**Figure 4.**
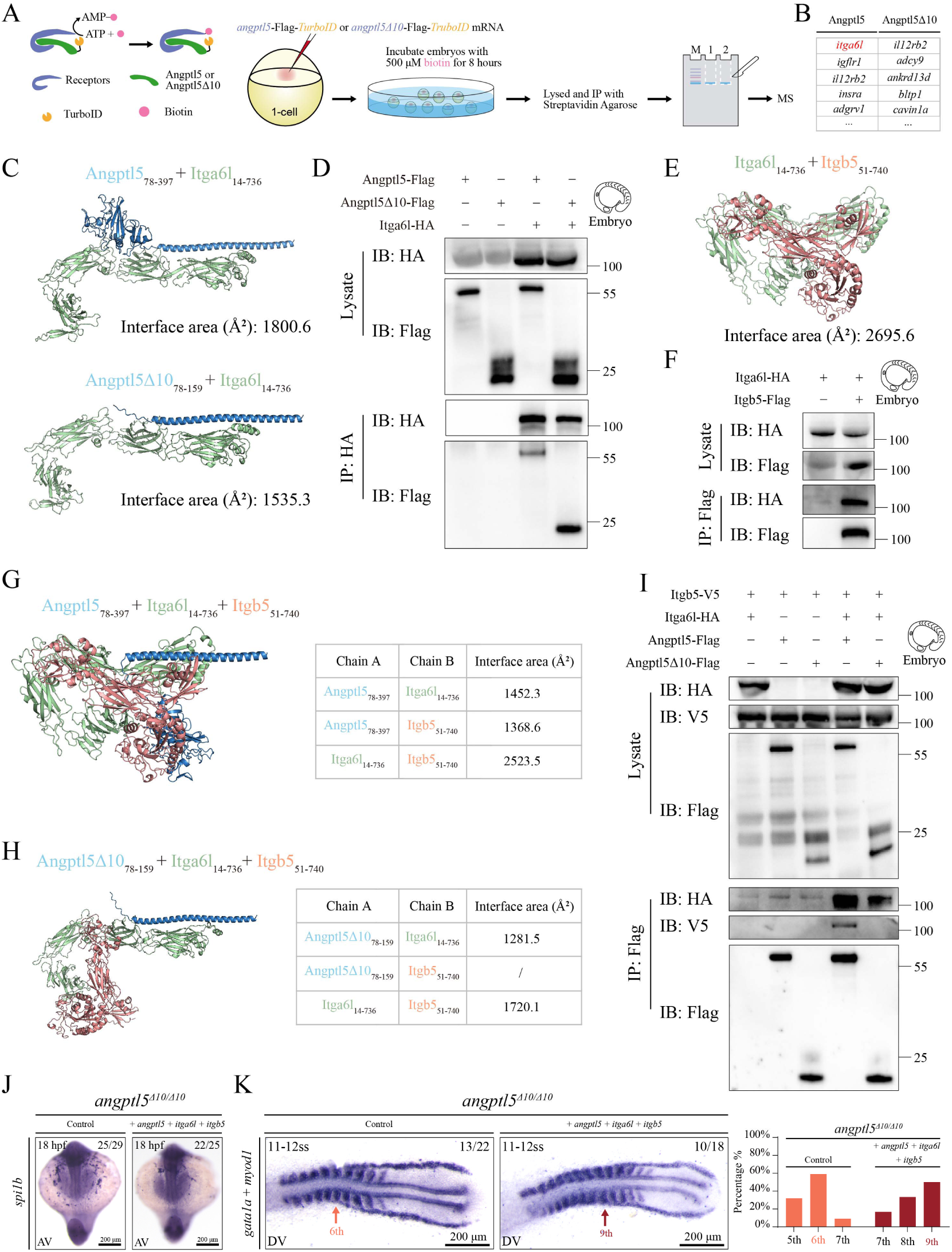
Angptl5 physically interacts with Integrin α6l and β5. (A-B) Candidate receptors of Angptl5 as assessed using immunoprecipitation-mass spectrometry (IP-MS). Schematic workflow of experimental setup for IP-MS (A) and identified proteins in Angptl5 and Angptl5Δ10 embryos were listed (B). (C) Structural models of the Angptl5_78-397_-Itga6l_14-736_ and Angptl5Δ10_78-159_-Itga6l_14-736_ complexes. (D) Interaction of Angptl5-Flag/Angptl5Δ10-Flag with Itga6l-HA in zebrafish embryos as assessed using co-immunoprecipitation (Co-IP). (E) Structural models of the Itga6l_14-736_-Itgb5_51-740_ complex. (F) Interaction of Itgb5-Flag with Itga6l-HA in zebrafish embryos as assessed using Co-IP. (G-H) Structural models of the Angptl5_78-397_-Itga6l_14-736_-Itgb5_51-740_ (G) and Angptl5Δ10_78-159_-Itga6l_14-736_-Itgb5_51-740_ (H) complexes. (I) Interaction of Angptl5-Flag/Angptl5Δ10-Flag, Itgb5-V5 and Itga6l-HA in zebrafish embryos as assessed using Co-IP. (J) WISH of *spi1b* in *angptl5^Δ10/Δ10^*embryos injected with *angptl5 + itga6l + itgb5* mRNA at the 1-cell stage. Uninjected embryos were used as control. (K) WISH of *gata1a* and *myod1* in flat-mounted *angptl5^Δ10/Δ10^* embryos injected with *angptl5 + itga6l + itgb5* mRNA at the 1-cell stage. Statistics are shown on the right. All structures were modelled using AlphaFold3. AV, anterior view; DV, dorsal view.

Comparative proteomic profiling of WT versus mutant samples, we identified integrin α6-like (Itga6l) as a potential candidate, based on its specific interaction with WT Angptl5 but not the Angptl5Δ10 mutant (Figure 4B, Dataset S1). This aligns with established literatures demonstrating that ANGPTL family members engage various integrins-conserved transmembrane proteins that regulate biochemical signaling and mechanotransduction via force-dependent conformational changes (13, 29). Structural modeling of the Angtpl5-Itga6l complex using AlphaFold3-Multimer also revealed a considerable interface area for the WT Angptl5 (Figure 4C, up) and co-immunoprecipitation (Co-IP) assay confirmed that Angptl5 can physically associate with Itga6l (Figure 4D). Notably, overexpression of *angptl5* mRNA in *angptl5^Δ10/Δ10^*embryos effectively suppressed the pathological expansion of *spi1b*^+^ myeloid progenitors, but this rescue phenotype was abrogated by co-injection with itga6l morpholino (MO), indicating that Angptl5-mediated regulation of primitive hematopoiesis is dependent on Itga6l function (Supplemental Figure 8D). Conversely, the Angptl5Δ10 exhibited a reduction interface area, indicating a reduced binding affinity for the truncated Angptl5 (Figure 4C, down). However, to our surprise, the co-immunoprecipitation (Co-IP) assay showed that both Angptl5 and Angptl5Δ10 can physically associate with Itga6l (Figure 4D). These finds suggests that the loss-of-function phenotype cannot be attributed solely to disrupted binding of Angptl5 to Itga6l.

Functional integrin signaling requires the formation of α/β heterodimeric complexes at the cell membrane (30). We thus speculate that the mutation of Angptl5 may disrupted the formation of this potential complex. To identify the β subunit partner for Itga6l in this context, we systematically evaluated potential candidates. Bioinformatic screening of existing transcriptome datasets revealed differential expression profiles across 12 integrin β subunit genes in zebrafish (31). Among these, five subunits (*itgb1a*, *itgb1b*, *itgb4*, *itgb5*, *itgb6*) showed detectable expression during primitive hematopoiesis (Supplemental Figure 9A). Thus, we performed structural modeling of the complexes formed between Itga6l and each individual β subunit. The Itga6l-Itgb4 and Itga6l-Itgb5 complexes exhibited the two largest interfacial contact areas among all Itga6l-β subunit complexes (Figure 4E, Supplemental Figure 9B-F). However, the Angptl5-Itga6l-Itgb4 ternary complex failed to maintain structural stability due to incomplete pairwise interaction interfaces (Supplemental Figure 9G). Instead, structural modeling demonstrated that Itgb5 engages in an extensive interaction with Itga6l, burying a surface area of 2695.6 Å² at their interface (Figure 4E), and this is further confirmed biochemically by Co-IP (Figure 4F). WISH analysis showed that both *itga6l* and *itgb5* are expressed during early development (Supplemental Figure 9H).

We therefore characterized the putative Angptl5-Itga6l-Itgb5 ternary complex. Structural modeling revealed that WT Angptl5 formed stable ternary interactions through extensive interfacial contact surfaces, whereas the Angptl5Δ10 mutant exhibited disrupted binding interfaces with insufficient pairwise interaction networks (Figure 4G-H). Intriguingly, Co-IP assays demonstrated that Angptl5 cannot bind Itgb5 directly in the absence of Itga6l (Figure 4I). However, when Itga6l is present, WT Angptl5, but not the Δ10 mutant, mediates formation of a stable complex with Itgb5 (Figure 4I). These findings demonstrate that Angptl5 facilitates assembly of Itga6l-Itgb5 heterodimers, and that this molecular activity is abolished in Angptl5Δ10 mutant.

To elucidate the functional role of the Angptl5/Integrin α6lβ5 complex in zebrafish primitive hematopoiesis, we performed gain-of-function experiments in WT and the mutant embryos. In WT embryos, either *angptl5* mRNA injection or co-injection of *itga6l* and *itgb5* mRNAs effectively suppressed *spi1b* expression, while *angptl5Δ10* mRNA had no significant effect (Supplemental Figure 9J). Notably, simultaneous overexpression of all three components (*angptl5*, *itga6l*, and *itgb5*) produced a further reduction in *spi1b* expression. Most importantly, in *angptl5^Δ10/Δ10^* mutants, this triple overexpression (*angptl5* + *itga6l* + *itgb5*) significantly reduced both the myeloid and erythroid hyperplasia (Figure 4J-K).

Collectively, these results suggest that integrin α6lβ5 acts as the cognate receptor for Angptl5 in hematopoietic regulation, and that the Angptl5-mediated suppression of primitive hematopoietic progenitor cells is mechanistically dependent on integrin α6lβ5 signaling.

### ERK signaling is required for Angptl5-mediated regulation of primitive hematopoiesis

Integrin engagement by extracellular ANGPTL proteins typically activates intracellular signaling cascades, including FAK, NF-κB, and MAPK/ERK pathways, which regulate diverse cellular processes such as motility, proliferation, and transcriptional regulation (32–34). To delineate the downstream effectors of Angptl5/α6lβ5 signaling in hematopoietic development, we systematically inhibited key pathways using small-molecule inhibitors from the shield stage onward, including: Defactinib (FAK), BAY 11-7082 (NF-κB), Adezmapimod (p38 MAPK), and Mirdametinib (ERK). Notably, FAK inhibition showed no significant effect on *spi1b* expression in the RBI, suggesting minimal involvement in primitive myeloid regulation (Supplemental Figure 10A). And NF-κB blockade downregulated *spi1b* expression, revealing its role in promoting myeloid expansion (Supplemental Figure 10B). Interestingly, MAPK or ERK inhibition significantly upregulated *spi1b* expression, and reduced *cyp26a1* expression, phenocopying *angptl5* loss-of-function mutants (Figure 5A, Supplemental Figure 10C). Most notably, pharmacological activation of ERK signaling (C16-PAF) effectively rescued the *spi1b+* myeloid hyperplasia in *angptl5^Δ10/Δ10^* mutants (Figure 5B).

**Figure 5.**
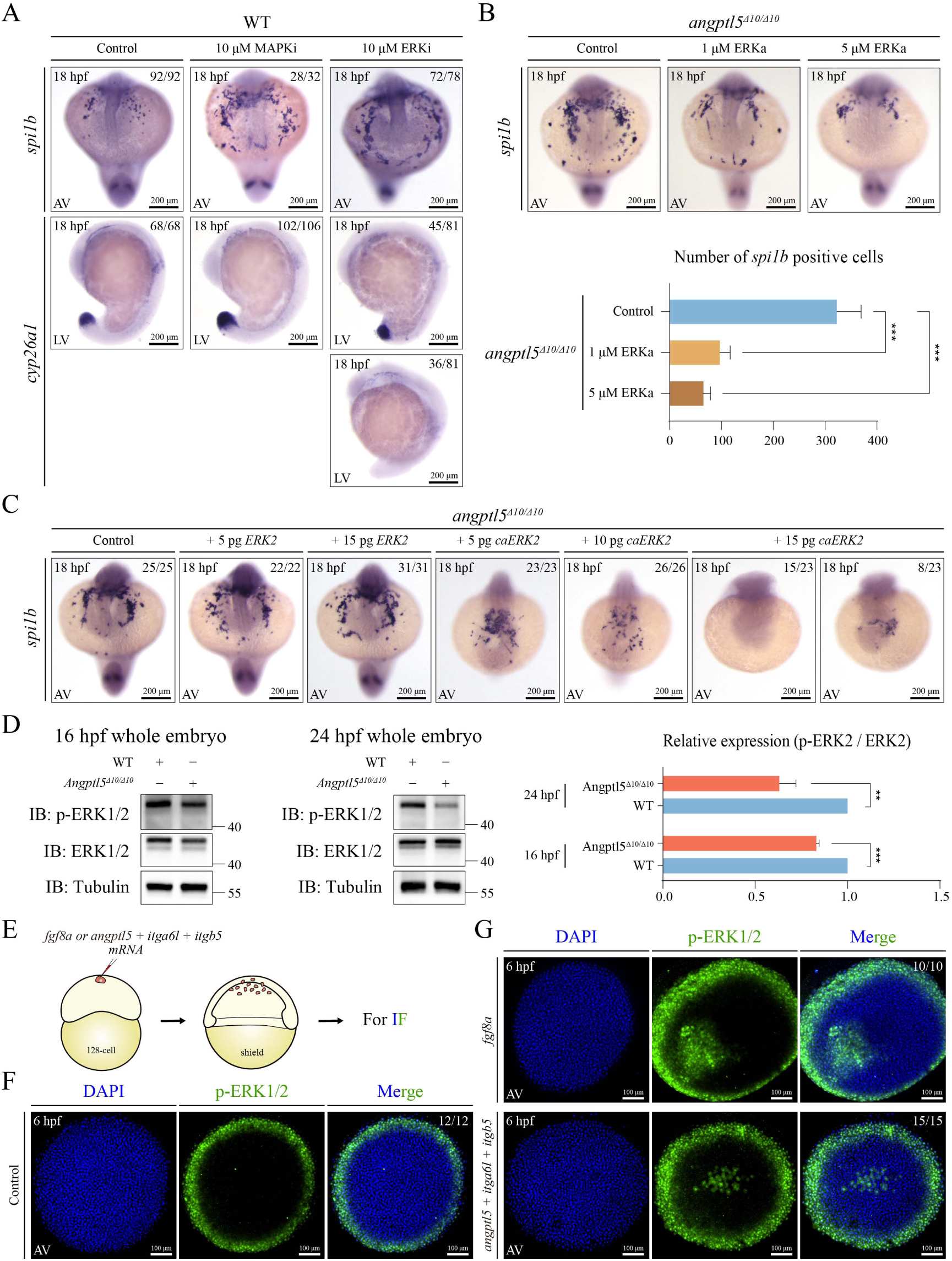
Angptl5-Integrin α6lβ5 functions through the ERK signaling pathway. (A) WISH of *spi1b* and *cyp26a1* in WT embryos treated with MAPK inhibitor or ERK inhibitor from the shield stage to the 18-somite stage. Untreated embryos were used as control. (B) WISH of *spi1b* in *angptl5^Δ10/Δ10^*embryos treated with ERK activator from the shield stage to the 18-somite stage. Untreated embryos were used as control. Statistics are shown on below. Data presented as the mean ± SD. Statistical significance: ***P < 0.001. (C) WISH of *spi1b* in *angptl5^Δ10/Δ10^* embryos injected with *ERK2* or *caERK2* mRNA at the 1-cell stage. Uninjected embryos were used as control. (D) Expression of phosphorylated ERK1/2 (p-ERK1/2) in WT and *angptl5^Δ10/Δ10^* embryos as assessed using western blot. Statistics are shown on the right. Three independent biological replicates were used. Statistical significance: **P < 0.01, ***P < 0.001. (E-G) Schematic diagram of experimental setup (E) for immunofluorescence (IF) of p-ERK1/2 in WT and *angptl5^Δ10/Δ10^* embryos. *Fgf8a* mRNA or *angptl5 + itga6l + itgb5* mRNA (G) were injected into one blastomere on the animal pole at the 128-cell stage and then imaged at the sphere stage. Uninjected embryos (F) were used as control. AV, anterior view (A-C), animal view (F-G); LV, lateral view.

The above results suggest that *angptl5* may regulate primitive hematopoiesis through the ERK signaling pathway. To further validate this hypothesis, we generated WT ERK2 and a constitutively activated ERK2 mutant (caERK2, L84P/S162D/D330N) (35) (Supplemental Figure 10D), and injected mRNA of these two versions to rescue the phenotype resulting from *angptl5* loss-of-function, respectively. While WT *ERK2* had no obvious effect on *spi1b* expression, 10 pg of *caERK2* rescued *spi1b* expression in *angptl5^Δ10/Δ10^* embryos to levels comparable to the WT, and higher doses of *caERK2* abolished *spi1b* expression in most *angptl5^Δ10/Δ10^* embryos and in all WT embryos (Figure 5C, Supplemental Figure 10E). Consistently, the ERK signaling activity was significantly decreased in *angptl^5Δ10/Δ10^* embryos compared to WT, as indicated by Western blot and immunofluorescence (IF) of phosphorylated ERK1/2 (pERK1/2) (Figure 5D, Supplemental Figure 10F).

Next, we investigated whether the Angptl5/Integrin α6lβ5 complex is sufficient to active ERK signaling. We found that overexpression of *angptl5* alone, co-overexpression of *itga6l* and *itgb5*, or combined overexpression of *angptl5* with *itga6l* and *itgb5* enhanced ERK signaling (Supplemental Figure 10G). To further validate this, we co-injected *angptl5*, *itga6l*, and *itgb5* mRNA into one blastomere of the animal pole at the 128-cell stage, and used *fgf8a*, a known canonical ERK activator (36) as a positive control (Figure 5E). Strikingly, combinatorial overexpression of *angptl5*, *itga6l*, and *itgb5* triggered ectopic ERK activation, similar to the activation pattern observed with overexpression of *fgf8a* (Figure 5F-G).

In summary, these findings delineate a pathway in which Angptl5 engages integrin α6lβ5 to activate ERK signaling, a process required for the regulation of primitive hematopoiesis by Angptl5.

### Angptl5/Integrin α6lβ5 potentiates RA activity via ERK-dependent *dhrs9* transcription

Building on our previous characterization of RA signaling in Angptl5-dependent hematopoiesis, we investigated whether the Angptl5/Integrin α6lβ5-ERK axis can directly modulates RA activity. Using *cyp26a1* as a readout of RA pathway activation, we found that ERK inhibition significantly reduced the *cyp26a1* expression in the WT embryos, while ERK activation drastically enhanced *cyp26a1* expression. Interestingly, triple overexpression of *angptl5*, *itga6l*, and *itgb5* at one cell stage also dramatically increase the *cyp26a1* expression. Importantly, this Angptl5/Integrin-mediated *cyp26a1* induction was completely abolished by ERK inhibition, demonstrating that the Angptl5/integrin complex enhances RA signaling through ERK-dependent mechanisms (Figure 6A).

**Figure 6.**
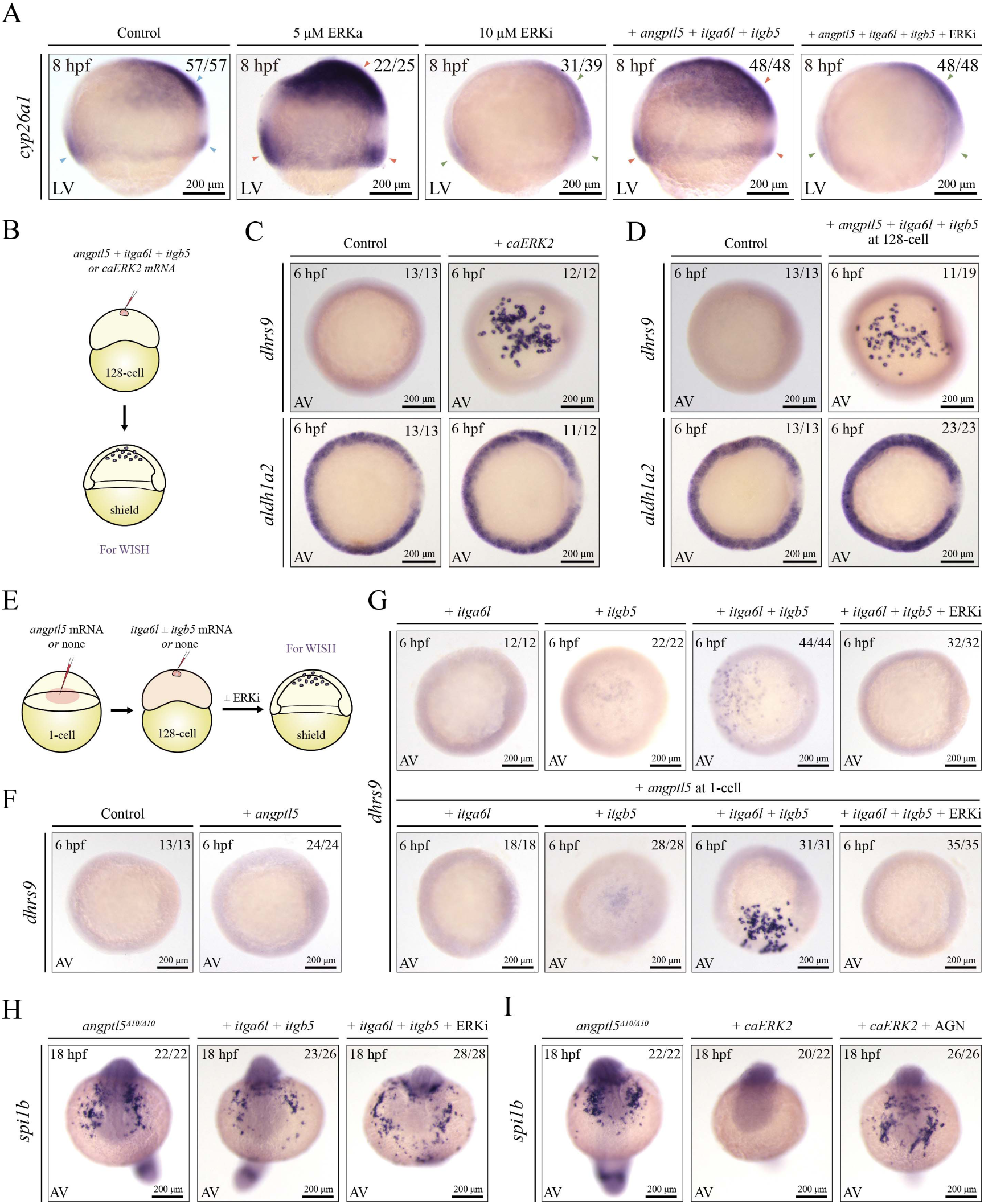
The Angptl5-Integrin α6lβ5-ERK-Dhrs9-RA signaling cascade regulates primitive hematopoiesis. (A) WISH of *cyp26a1* in WT embryos. Embryos were treated with ERK activator or ERK inhibitor from the shield stage, or injected with *angptl5* + *itga6l* + *itgb5* mRNA at the 1-cell stage. Untreated embryos were used as control. (B-D) Schematic diagram of experimental setup (B) for WISH of *dhrs9* and *aldh1a2* in WT embryos. *caERK2* (C) or *angptl5* + *itga6l* + *itgb5* (D) mRNA injected into one blastomere on the animal pole at the 128-cell stage and then detected at the sphere stage. (E-G) Schematic diagram of experimental setup (E) for WISH of *dhrs9*. WT embryos were first injected with *angptl5* mRNA at the 1-cell stage. Subsequently, *itga6l ± itgb5* mRNA were injected into one blastomere on the animal pole at the 128-cell stage. Embryos were then continuously treated with or without ERK inhibitor until the shield stage (G). Uninjected embryos and only *angptl5* mRNA injected embryos (F) were used as control. (H-I) WISH of *spi1b* in *angptl5^Δ10/Δ10^* embryos. Embryos were injected with *itga6l + itgb5* mRNA at the 1-cell stage and treated with or without ERK inhibitor from the shield stage to the 18-somite stage (H), or injected with *caERK*2 mRNA at the 1-cell stage and treated with or without RA receptor antagonist AGN 193109 (I). Uninjected embryos were used as control. LV, lateral view; AV, animal view (C-G), anterior view (H-I).

To further explore the mechanism by which the Angptl5/Integrin α6lβ5-ERK axis regulates RA activity, we injected *caERK2* mRNA or *angptl5*, *itga6l*, and *itgb5* mRNA together into one blastomere at the 128-cell stage, followed by analysis of the expression of RA synthases *aldh1a2* and *dhrs9* (Figure 6B). Interestingly, both *caERK2* and triple overexpression of *angptl5*, *Itga6l*, and *Itgb5* mRNA induced *dhrs9* expression at the animal pole, but not *aldh1a2* (Figure 6C-D). To further dissect the activating effects of Angptl5, Itga6l, Itgb5, and ERK on *dhrs9* transcription, we performed a two-stage microinjection in zebrafish embryos (Figure 6E). Our results showed that neither ectopic expression of *angptl5* nor *itga6l* alone induced *dhrs9* expression. Intriguingly, individual administration of *itgb5* mRNA, or its co-delivery with *angptl5* mRNA, slightly elicited *dhrs9* induction. Notably, co-injection of *itga6l* and *itgb5* mRNA provoked moderate transcriptional activation of *dhrs9*, whereas simultaneous overexpression of *angptl5* dramatically boosted this activation. Strikingly, this synergistic transcriptional response was completely ERK signaling-dependent, as evidenced by its complete abrogation upon pharmacological ERK inhibition (Figure 6F-G). These findings delineate a mechanistic cascade wherein Angptl5, together with Itga6l/Itgb5, activates ERK-dependent signaling to drive *dhrs9* transcription and subsequently modulate RA activity, which further supported by that co-overexpression of *angptl5*, *itga6l* and *itgb5* significantly potentiated RA signaling in *angptl5^Δ10/Δ10^*embryos; and this potentiation was markedly attenuated by ERK inhibition (Supplemental Figure 11A).

Next, we validated the regulatory role of the Angptl5/Integrin α6lβ5-ERK-RA axis in primitive hematopoiesis. Co-injection of *itga6l* and *itgb5* mRNA into *angptl5^Δ10/Δ10^*embryos partially attenuated myeloid hyperplasia, but this suppression was reversed upon ERK inhibitor treatment, revealing that the regulatory function of integrin α6lβ5 in primitive hematopoiesis is mechanistically reliant on ERK signaling activation (Figure 6G). In contrast, *caERK2* overexpression completely abolished *spi1b* expression in *angptl5^Δ10/Δ10^* embryos, but this inhibitory effect was counteracted by RA inhibitor treatment, suggesting that ERK signaling-mediated regulation of primitive hematopoiesis requires RA activity (Figure 6H).

Taken together, these findings establish Angptl5 as a critical regulator of zebrafish primitive hematopoiesis, operating through the Angptl5/Integrin α6lβ5-ERK-RA signaling axis to suppress excessive expansion of hematopoietic progenitor cells.

## Discussion

Our study establishes Angptl5 as a new regulator of primitive hematopoiesis in zebrafish, operating through a previously unrecognized Angptl5-Integrin α6lβ5-ERK-retinoic acid signaling axis. By combining genetic, biochemical and structural modeling approaches, we demonstrate that Angptl5-mediated suppression of hematopoietic progenitor expansion requires sequential activation of integrin α6lβ5 heterodimerization, ERK signaling, and subsequent Dhrs9-dependent RA production (Figure 7).

**Figure 7.**
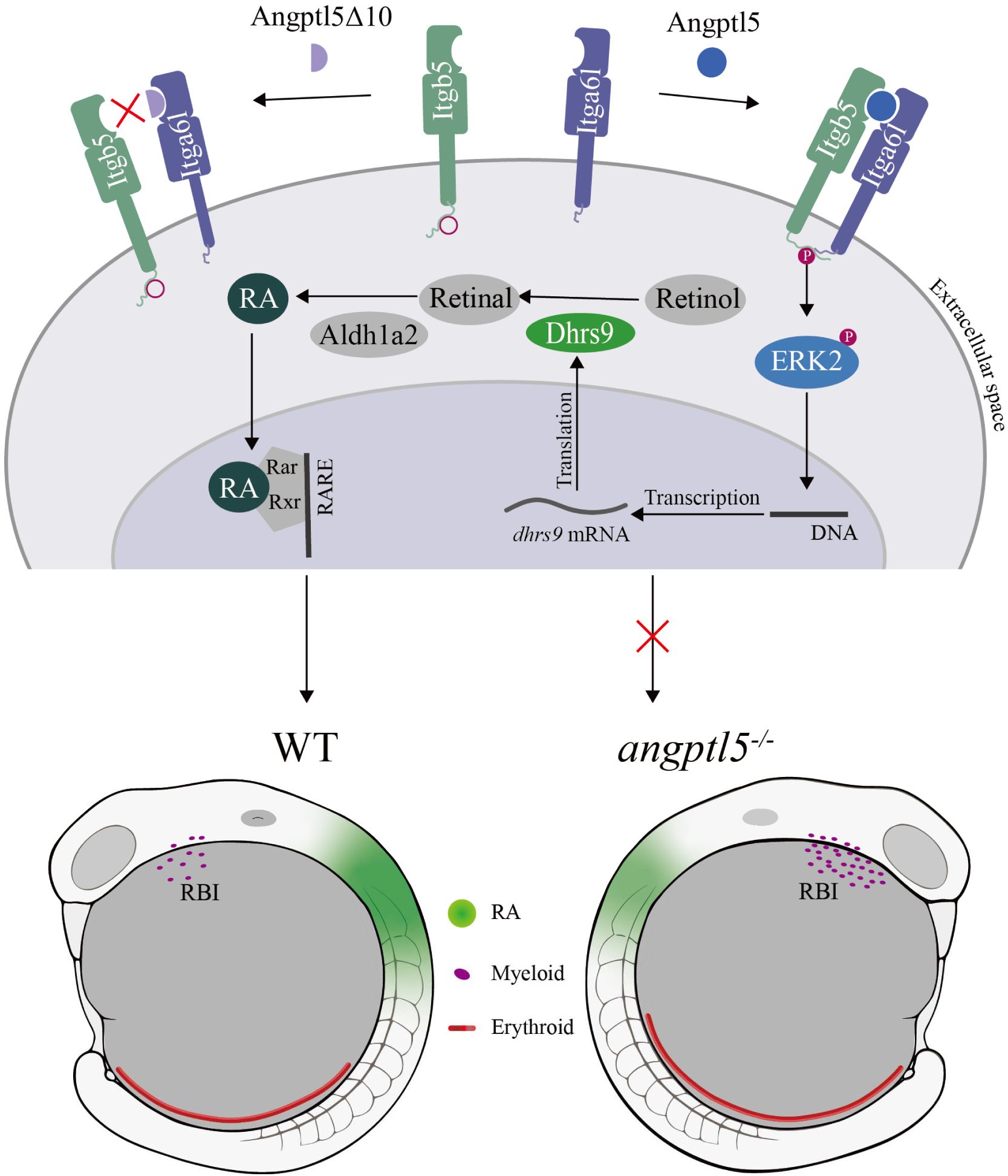
Working model of Angptl5-mediated regulation of primitive hematopoiesis in zebrafish. During primitive hematopoiesis, Angptl5 (but not the truncated Angptl5Δ10) specifically promotes the assembly of Itga6l and Itgb5 subunits into functional integrin α6lβ5 heterodimers, thereby activating downstream ERK signaling pathways. Mechanistically, activated ERK2 directly induces dhrs9 expression through transcriptional activation, which potentiates RA signaling to suppress excessive expansion of myeloid and erythroid progenitors in primary hematopoiesis.

Members of the Angiopoietin-like (Angptl) protein family play crucial roles in various physiological and pathological processes, including metabolism (37, 38), inflammation (13, 39) and cancer (40). Several members of this family, including Angptl5, have been reported to *ex vivo* positively regulate the maintenance and proliferation of stem cells, including hematopoietic stem cells (HSCs) (19, 41, 42). However, in our study, we observed that the loss of Angptl5 did not lead to significant changes in primitive HSCs. This may be due to the redundancy or compensatory mechanisms provided by other Angptl family members in vivo.

Interestingly, the loss of Angptl5 led to the simultaneous expansion of myeloid cells at the anterior of the body axis and erythroid cells at the posterior, despite their established mutual antagonism (43, 44). This suggests that the hematopoietic microenvironment plays a specific and critical role in maintaining the proliferative capacity of hematopoietic progenitors at different stages. This also raises an important question: despite both anterior and posterior hematopoietic progenitors originating from the same primitive HSC (45), what signals in the anterior and posterior microenvironments induce their distinct differentiation into myeloid and erythroid progenitors, respectively? How does the Angptl5-RA axis interact with these key signals? These questions provide an exciting avenue for future research.

Our mechanistic studies identify ERK as a central hub in Angptl5-dependent hematopoietic control, promoting retinoic acid (RA) synthesis through transcriptional activation of dhrs9. Notably, this forms a potential feedback loop, as previous studies show that RA can rapidly activate ERK via non-canonical, receptor-independent mechanisms (46, 47). This bidirectional interaction between RA and ERK suggests a tightly regulated circuit with possible roles in fine-tuning hematopoietic processes. Further investigation is warranted to explore how this circuit functions across developmental stages, tissues, and in both normal and disease contexts.

In summary, we identify Angptl5 as a critical rheostat of primitive hematopoiesis, bridging integrin-mediated signaling (Itga6l/Itgb5) with RA metabolic regulation (dhrs9). This axis ensures precise control of myeloid-erythroid progenitor expansion during embryogenesis, with perturbations leading to hematopoietic hyperplasia. These findings not only expand the known functions of ANGPTL family proteins beyond angiogenesis and metabolism but also reveal an intricate signaling cascade that coordinates hematopoietic homeostasis during embryonic development. Furthermore, we uncover a novel crosstalk between ECM sensing and RA gradients—a mechanism potentially conserved across vertebrates. In addition, targeting the Angptl5-integrin-ERK axis may offer therapeutic avenues for disorders characterized by RA-ERK imbalance, such as chemotherapy-resistant AML (48).

## Materials and Methods

All fish maintenance were performed per the requirements of the “Regulation for the Use of Experimental Animals in Zhejiang Province”, with the approval by the Zhejiang University Animal Care and Use Committee. The protocol number is ZJU20220375. Specific experimental details about whole-mount in situ hybridization (WISH), morpholino knockdown, generation of *angptl5*^Δ10/Δ10^ and *angptl5*^Δ5/Δ5^ mutants, *Tg(angptl5:*EGFP) and *Tg(RARE:*EGFP), in situ hybridization chain reaction (HCR), immunofluorescence (IF), quantitative real-time PCR (qPCR), western blot (WB), co-immunoprecipitation (Co-IP), Single-cell sequencing (scRNA-seq), and the following bioinformatics analysis; are described in supporting information. The primers used in this study are listed in SI Appendix, Tables S1 (SI Appendix, Tables S1).

Other Materials and Methods details can be found in the Supporting Information.

## Data availability

Single-cell RNA sequencing datasets generated in this study are deposited in the Gene Expression Omnibus (GEO) repository under accession number GSE295891. Previously published datasets analyzed in this work are available through GEO under accession GSE223922 (16 hpf developmental stage) and GSE112294 (18 hpf and 24 hpf stages).

## Author Contributions

P.F.X., L.P.S. and Z.X.H. designed the research; J.M., D.H.Z., Y.H., T.C., Y.D., Y.Y.X., Y.F.L., L.X., Z.X.J., H.R.P., G.Q.Z, and Y.M.L. performed research; T.C. and Y.D. analyzed the scRNA-seq data; J.M., D.H.Z., P.F.X., L.P.S. and Z.X.H wrote the manuscript; P.F.X., L.P.S. and Z.X.H reviewed the manuscript.

## Acknowledgments

We would like to thank Yu Feng (Department of Biophysics, ZheJiang University), Ruixing Hu and Qing Lu (Bio-X Institutes, Key Laboratory for the Genetics of Developmental and Neuropsychiatric Disorders (Ministry of Education), Shanghai Jiao Tong University) for their expert guidance in protein-protein interaction structural prediction. We also thank the members of Laboratory of Development and Organogenesis (LDO) at Zhejiang University for helpful suggestions and discussions. We thank Ying-Niang Li from the zebrafish core facility at Zhejiang University School of Medicine for their technical support. This work was supported by grants from the National Natural Science Foundation of China (32470851, 32300688, 32160168, 32360214, 32300677), in part by the Guizhou Province’s Science and Technology Major Project (GCC[2023]034), and Chinese National Key Research and Development Project (2022YFA1103102, 2024YFA1803001).

**Supporting Information for**

## Materials and Methods

### Zebrafish husbandry and genetic strains

Zebrafish were raised on a 14-hours-light/10-hours-dark cycle at 28°C (1) and staged as described (2). The following lines were used in this study: wild-type (WT) AB strains, *Tg(angptl5:EGFP), Tg(RARE:EGFP)*, *Tg(kdrl:GFP)*, *angptl5*^Δ10/Δ10^ and *angptl5*^Δ5/Δ5^ mutants. All fish maintenance were performed per the requirements of the “Regulation for the Use of Experimental Animals in Zhejiang Province”, with the approval by the Zhejiang University Animal Care and Use Committee.

### Whole-mount in situ hybridization (WISH) and histological sections

WISH of zebrafish was performed essentially as described previously (3). Probe templates including *tal1*, *spi1b*, *gata1a*, *mpx*, *lyz*, *mpeg1.1*, *myod1*, *aldh1a2*, *dhrs9*, *cyp26a1*, *angptl1-7*, *etsrp* and *kdrl* have been generated from zebrafish cDNA and clones inserted into the pEASY-blunt-zero vector (Transgen). Antisense RNA probes were labeled with DIG RNA labeling Mix (Roche) by transcription using T7 or T3 polymerase (Promega) *in vitro*.

For histological sections, embryos were washed with PBS following WISH and embedded in OCT compound (optimal cutting temperature compound, Sakura). Cryosections of 20 μm thickness were prepared along the anterior–posterior axis using an NX50 Cryostat Microtome (Thermo Scientific), and images were captured with an ECLIPSE Ni microscope.

### Generation of Tg(*angptl5*:EGFP) and Tg(*RARE*:EGFP) transgenic lines

The *angptl5:EGFP* plasmid was generated by cloning the *angptl5* promoter by cloning the upstream 867 bp region of *angptl5* gene and EGFP fragment into the Tol2 backbone using a seamless cloning kit (Vazyme). For the *RARE:EGFP* plasmid, the EF1α enhancer-promoter in the PT2AL200R150G transposon vector (4) was replaced with a trimerized *RARE* promoter sequence, which was amplified from the pGL3-RARE-luciferase plasmid (Addgene). Both plasmids were co-injected with *Tol2* mRNA into zebrafish embryos at the 1-cell stage to generate the *Tg(angptl5:EGFP)* and *Tg(RARE:EGFP)* transgenic lines, respectively.

### *In situ* hybridization chain reaction (HCR)

HCR staining was performed as described previously (5). Briefly, zebrafish embryos were fixed with 4% (w/v) paraformaldehyde in DEPC-treated PBS at 4°C overnight. Samples underwent proteinase K (MCE) treatment before incubation of probe. Samples were incubated with 2 pmol of each HCR probe sets in 100 μL probe hybridization buffer at 37°C overnight. Hybridization was terminated by washing samples repeatedly with probe wash buffer at room temperature. For amplification, the probes were conjugated with fluorescent HCR amplifiers in amplification buffer for 4 h at room temperature in the dark. The reaction was stopped by washing several times with 5× SSCT (DAPI co-staining was achieved by adding 0.1 μg/mL DAPI). Fluorescent hairpins, buffers and DNA probes were purchased from Molecular Technologies. Fluorescent signals were detected using an OLYMPUS CSU-W1 confocal microscope.

### Generation of *angptl5^Δ10/Δ10^* and *angptl5*^Δ5/Δ5^ mutants using CRISPR/Cas9

To generate the *angptl5^Δ10/Δ10^* and *angptl5^Δ5/Δ5^*mutants, we synthesized gRNAs targeting the fourth exon and third exon of the zebrafish *angptl5* gene respectively. These procedures were performed as previously described (6). The guide RNAs (Supplementary Table 1), targeting zebrafish *angptl5* gene, were designed using CHOPCHOP (http://chopchop.cbu.uib.no) and synthesized using the T7 High Yield RNA Transcription Kit (Vazyme). The SpyCas9 protein (NEB) and *angptl5*-targeting gRNA were co-injected into wild type embryos at the 1-cell stage. The *angptl5* mutant lines were identified in the F1 generation by analyzing the PCR products using the primer pair listed in Supplementary Table 1.

### Immunofluorescence

Whole-mount immunofluorescence staining was performed as described previously (7) using the following primary antibody: Phospho-Erk1 (Thr202/Tyr204)/Erk2 (Thr185/Tyr187) rabbit monoclonal antibody (Beyotime, 1:100), and Alexa Fluor 488-conjugated donkey anti-rabbit antibody (Invitrogen, 1:500 dilution) used as the secondary antibody. Embryos were photographed using an OLYMPUS CSU-W1 confocal microscope.

### Preparing samples for sequencing and pro-processing single cell RNA-seq data

The *angptl5^Δ10/Δ10^* embryos were raised in 0.3× Danieau Buffer and harvested at 16 hpf. Collected samples were digested with trypsin into single cell suspension and set to Novogene for single cell sequencing.

### Pro-processing single cell RNA-seq data

The raw Illumina sequencing reads were aligned to the zebrafish reference genome (GRCz11) through 10× Genomics CellRanger pipeline v3.0.2 with default parameters. The *angptl5^Δ10/Δ10^* embryos yielded 7,350 cells, and the generated expression matrix was further processed by Seurat v5.1.0 (8). Low-quality cells with fewer than 200 detected genes, more than 7000 detected molecules, or greater than 15% mitochoridal expression were filtered out. The single cell RNA-seq dataset of 16 hpf wild-type (WT) embryos was obtained from the pubished work (9). The single cell RNA-seq datasets of WT and *angptl5^Δ10/Δ10^*mutant embryos were merged for the further analyses. The NormalizeData function with default settings was used to normalized the raw counts. The FindVariableFeatures function was used to identify the highly variable genes. The normalized data was then scaled with the through ScaleData function. Principle component analysis (PCA) and UMAP were employed to reduce the dimensionality of the data.

### Integrative single cell RNA-seq analysis

The merged datasets were integrated using IntegrateLayers function with the canonical correlation analysis (CCA) integration method, utilizing shared principal components across two conditions. The layers of two conditions were re-joined after integration through the JoinLayers function. Spot clusters were obtained using the FindNeighbors and FindClusters function with the first 30 dimensions and the resolution set to 0.8. Non-linear reduction was run using RunUMAP function with the first 30 dimensions. Marker genes for each cluster were identified using the FindAllMarkers function, and each cluster was then annotated based on the expression of these marker genes.

Differential abundance analysis was conducted within neighbourhoods using the negative Binomial generalized linear model (GLM), with p-values corrected using the Spatial FDR method. A cell type label was assigned to each neighbourhood by identifying the most abundant cell type among the cells within that neighbourhood. A beeswarm plot was used to visualize the distribution of differential abundance fold changes among different cell types.

### Chemical treatment

In general, embryos were treated with drugs from the shield stage until collection at the indicated stages. When injected at the 128-cell stage, embryos received drug treatment immediately post-injection with continuous exposure until specimen collection at the shield stage.

The drug was initially dissolved in DMSO to prepare a high-concentration stock solution, which was subsequently diluted with 0.3×Danieau buffer to the working concentration. Detailed pharmacological parameters and experimental concentrations are tabulated below: Retinoic acid (RA) (sigma), Aldehyde dehydrogenase inhibitors 4-diethylaminobenzaldehyde (DEAB) (MCE, 10 μM), Retinoic acid receptor (RARs) antagonists AGN 193109 (MCE, 20 μM), FAK inhibitor Defactinib (MCE, 2 μM /8 μM), MAPK inhibitor Adezmapimod (MCE, 10 μM), ERK inhibitor Mirdametinib (MCE, 10 μM), ERK activator C16-PAF (MCE, 1 μM /5 μM), NF-κB inhibitor BAY 11-7082 (MCE, 0.1 μg/mL, 0.4 μg/mL).

### Quantitative Real-time PCR

Total RNA was isolated from zebrafish embryos (50 embryos per sample) using RNA isolater Total RNA Extraction Reagent (Vazyme), followed by cDNA synthesis with HiScript III 1st Strand cDNA Synthesis Kit (+gDNA wiper) (Vazyme). Quantitative real-time PCR (qPCR) was conducted on a LightCycler 480 Instrument II system (Roche Diagnostics) with 2× Universal SYBR Green Fast qPCR Mix (ABclonal). The ΔΔCT method employed for data analysis, with normalization to 18S ribosomal RNA (18s rRNA) levels. Primer sequences with annealing temperatures are detailed in Supplementary Table 1.

### Morpholino injections

The *itga6l*-morpholinos were purchased from Gene Tools LLC, and sequence of morpholino is 5’-GCTCTCCTTTCTTCATCAGGTTCAT-3’. Zebrafish embryos at the one-cell stage were microinjected with 60 nM morpholino solution using standard procedures.

### Plasmid constructs

The full-length *angptl5* coding sequence was amplified and cloned into a pCS2(+) vector with the addition of Flag or Flag-TurboID tag in frame at the C terminus. Based on Flag or Flag-TurboID tagged Angptl5 plasmid, Angptl5Δ10 was constructed by deleting 10bp (base pair 458 to 467). Angptl5 and Angptl5Δ10 tagged with Flag or V5 was used for Immunoblotting (IB) and co-immunoprecipitation (Co-IP) in zebrafish embryos. Angptl5 and Angptl5Δ10 tagged with Flag-TurboID was used for Immunoprecipitation-Mass Spectrometry (IP-MS) in zebrafish embryos. For Co-IP analysis in zebrafish embryos, the *itga6l* and *itgb5* coding sequences were amplified and cloned into a pCS2(+) vector, with addition of HA, V5 or Flag tag in frame at the C terminus.

### Co-immunoprecipitation and mass spectrometry

Co-immunoprecipitation (Co-IP) analysis in zebrafish embryos was performed as described previously (10). Briefly, embryos were co-injected with the indicated mRNA as described in the main text at the 1-cell stage, and collected at 5-6 hpf. Then embryos were lysed with ice-cold lysis buffer [50 mM Tris at pH 7.5, 150 mM NaCl, 10% glycerol, 1% Triton X-100 and complete protease inhibitor (Roche)]. Lysates were centrifuged and the supernatants were then transferred to a Spin Column (Pierce). Samples were incubated with HA-Nanoab-Agarose (LabLEAD), Anti-Flag Affinity Gel (Bimake) or DYKDDDDK-Nanoab-Agarose (LabLEAD) at 4 °C overnight, followed by four washes with ice-cold wash buffer (50 mM Tris at pH 7.5, 150 mM NaCl, 1%Triton X-100 and complete protease inhibitor) four times before adding IgG Elution Buffer (Pierce).

Immunoprecipitation-mass spectrometry (IP-MS) was performed as described (11). Briefly, 800 embryos were lysed and subjected to immunoprecipitation with Streptavidin Agarose Resins (Pierce). Samples were separated by SDS-PAGE. The excised gels were analyzed by liquid chromatography-tandem mass spectrometry (LC-MS/MS) at Shanghai Applied Protein Technology Co., Ltd (APTBIO, China).

### Western blot analysis

Western blot analysis was performed as described previously (12). Briefly, embryos or HEK293T cells were lysed with ice-cold RIPA Complete Lysis Buffer (Beyotime). For HEK293T conditioned medium (CM) preparation, the plasmids were transfected using polyethylenimine (PEI, Polysciences) at 1:2.5 DNA ratio. At 24 h post-transfection, medium was replaced with serum-free DMEM. After 24 hours of incubation, CM samples were collected, centrifuged and concentrated 5 to 10-fold using centrifugal filters (Millipore). Proteins were subjected and separated on SDS-PAGE gel and transferred to PVDF membranes (Millipore). After blocking with non-fat milk (CST), membranes were incubated overnight with the following specific primary antibodies: ERK1/2 (Beyotime), Phospho-Erk1 (Thr202/Tyr204)/Erk2 (Thr185/Tyr187) (Beyotime), Beta Tubulin (Proteintech), Flag-Tag (Beyotime), HA-Tag (CST) and V5-Tag (Invitrogen). After four TBST washes, membranes were incubated with appropriate HRP-conjugated secondary antibodies. After incubation, membranes were subjected to five TBST washes, then visualized using BeyoECL Moon Kit (Beyotime) or BeyoECL Star Kit (Beyotime) and detected with gel image analysis system (Tanon).

### Statistical analysis

Raw image data were processed using ImageJ software. Significance was calculated using independent samples t-tests in GraphPad Prism. The statistical parameters were shown in the figures and described in the figure legends or in the main text.

---

**Figure S1.**
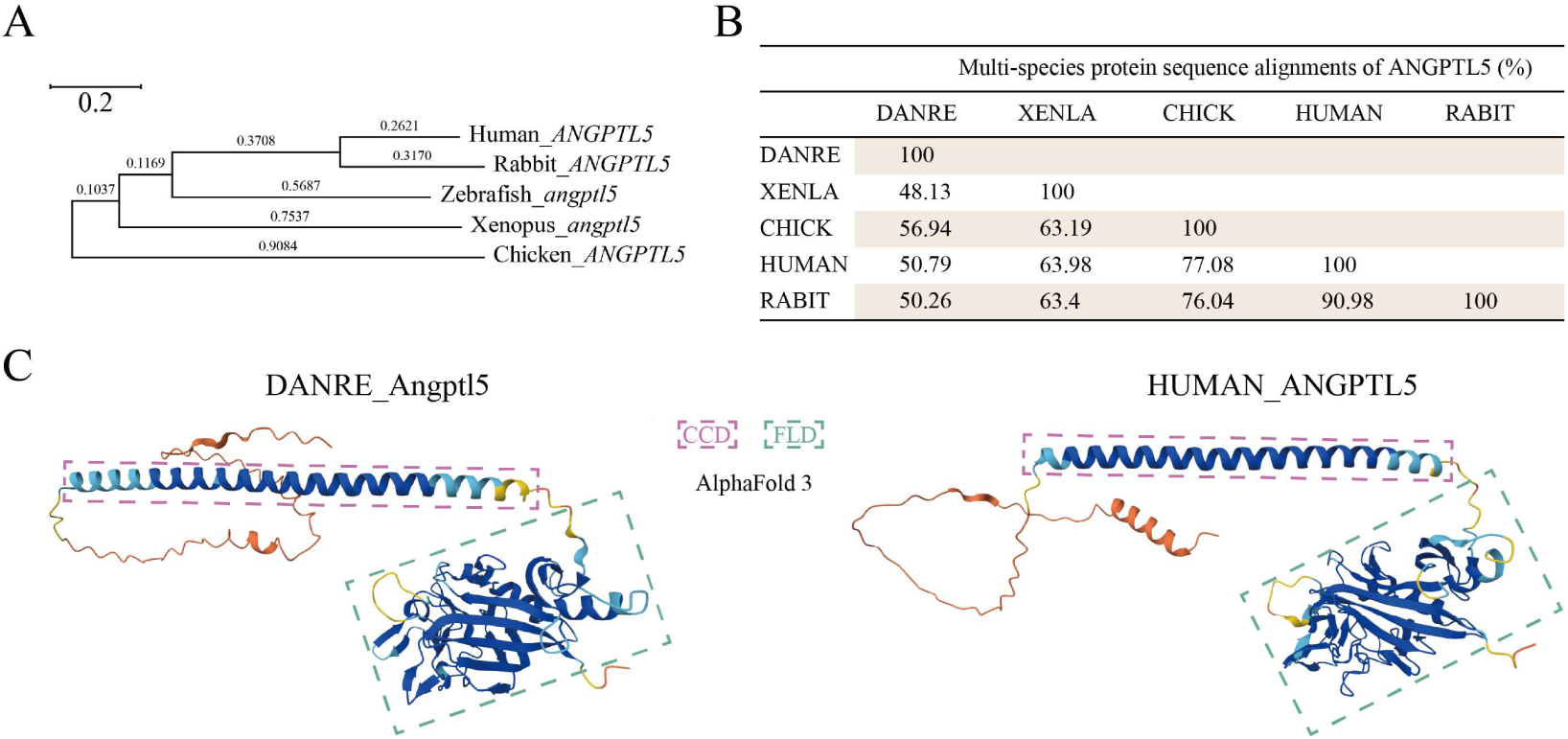
Cross-species evolutionary comparison of ANGPTL5. (A) Phylogenetic tree of *ANGPTL5* nucleotide sequence in human, rabbit, chick, *Xenopus* and zebrafish using the neighbor joining method. (B) ANGPTL5 protein sequence homology among different species. (C) Structural models of the zebrafish and human ANGPTL5 proteins were modelled using AlphaFold3. CCD, coiled-coil domain; FLD, fibrinogen-like domain.

**Figure S2.**
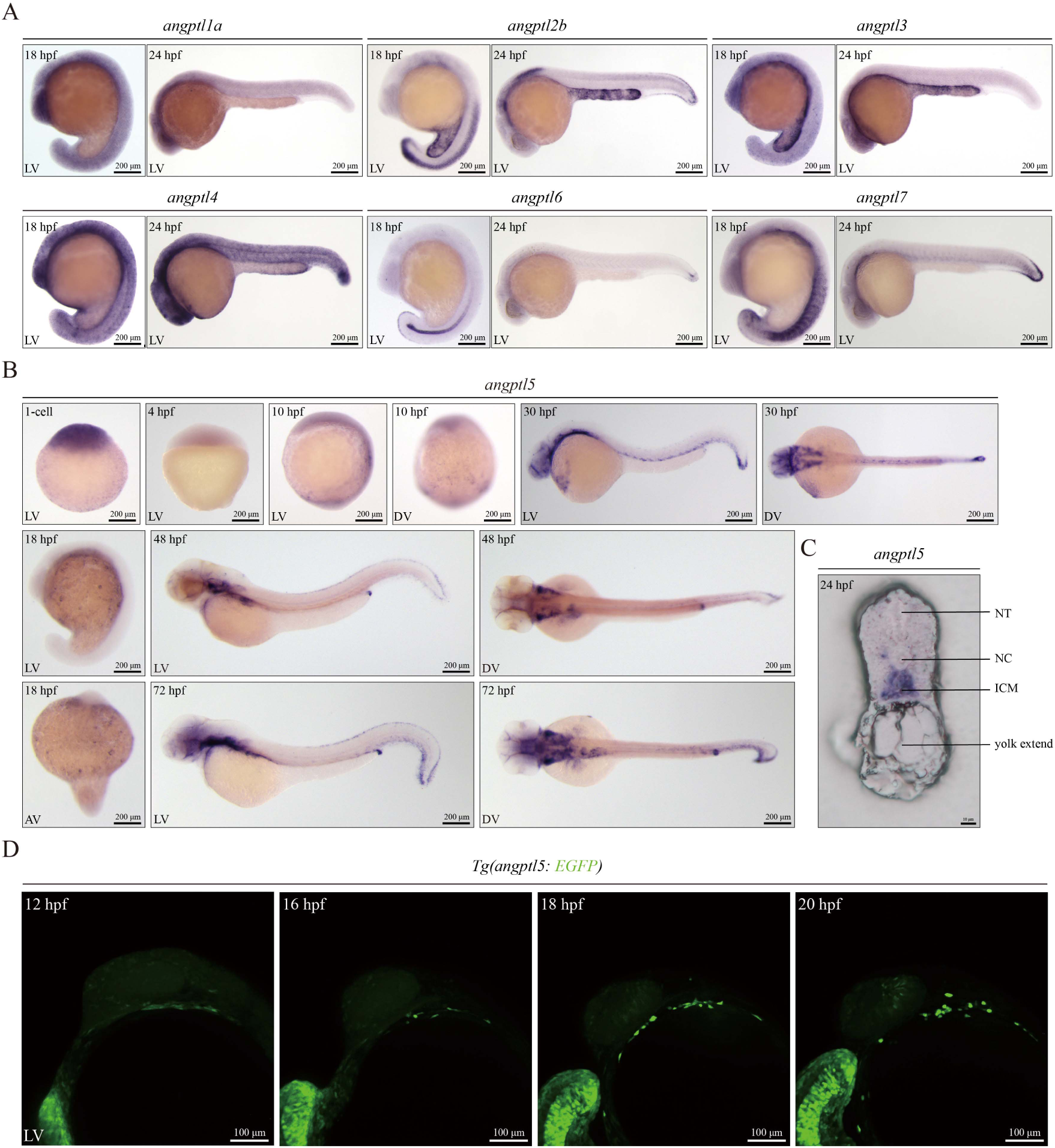
Expression patterns of *angptl* family members during zebrafish embryonic development. (A) WISH of *angptl* family members in zebrafish embryos. (B) WISH of *angptl5* during zebrafish embryonic development at the indicated stages. (C) Cryosection of the trunk region for 24 hpf zebrafish embryo using O.C.T. after WISH of *angptl5*. (D) Time-lapse confocal micrographs of the *angptl5* positive cell in *Tg(angptl5:*EGFP) embryos. Each imaging was performed for at least three independent replicates. LV, lateral view; DV, dorsal view; AV, anterior view. NT, Neural tube; NC, notochord; ICM, intermediate cell mass.

**Figure S3.**
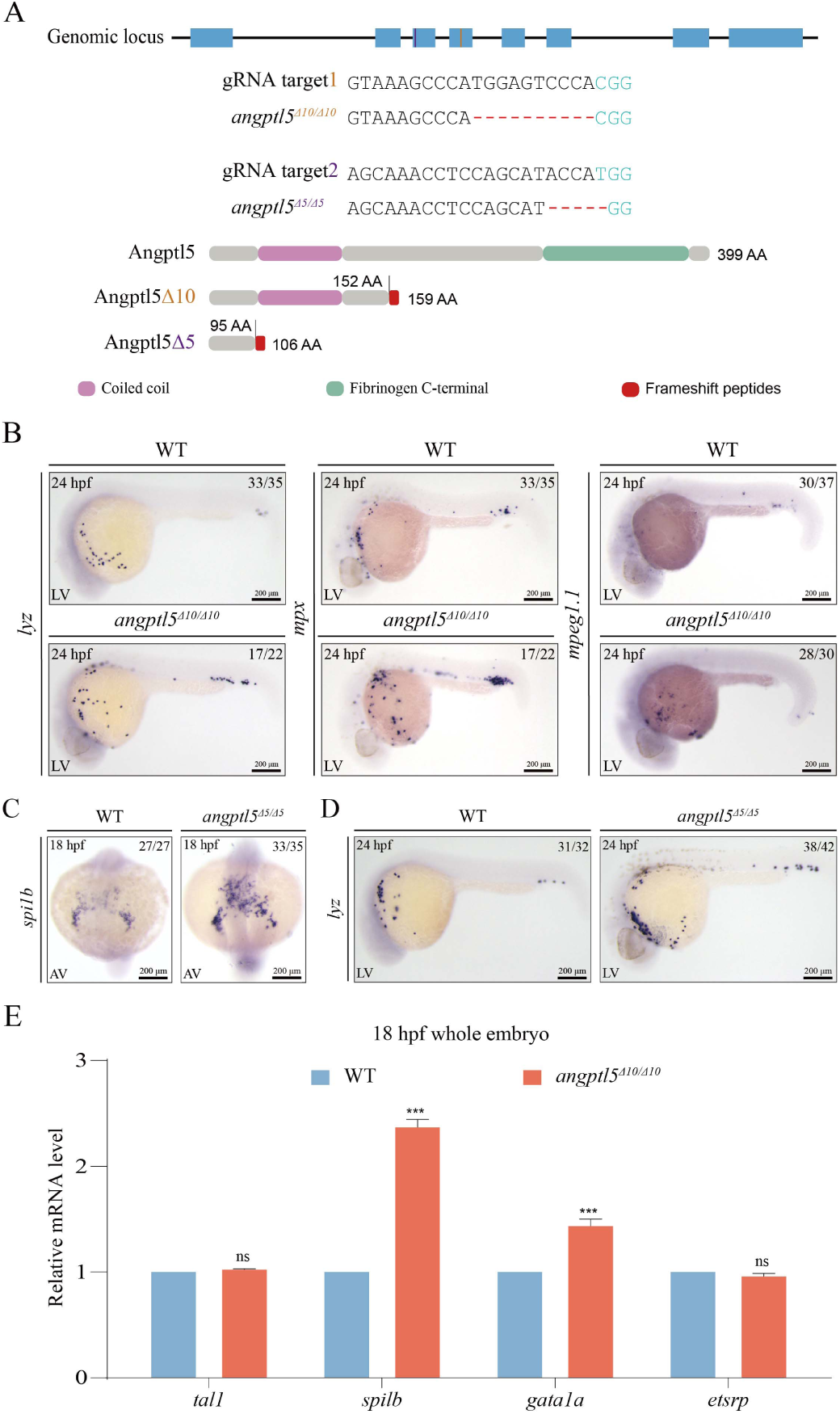
Comparison of primitive hematopoiesis between WT and *angptl5^Δ10/Δ10^* embryos. (A) Schematic diagram showing the positions of gRNA target site; the deletions of *angptl5* mutant lines generated using CRISPR/Cas9; and the predicted truncated Angptl5Δ10 and Angptl5Δ5 proteins (predicted domains taken from the Uniprot database). (B) WISH of neutrophil markers *lyz, mpx* and macrophage marker *mpeg1.1* in WT and *angptl5^Δ10/Δ10^*embryos. (C-D) WISH of *spi1b* (C) and *lyz* (D) in WT and *angptl5^Δ5/Δ5^*embryos. (E) qPCR shows the *tal1*, *spi1b*, *gata1a* and *etsrp* expression in WT and *angptl5^Δ10/Δ10^* embryos. Data presented as the mean ± SD. Three independent biological replicates were used. Statistical significance: ***P < 0.001. LV, lateral view; AV, anterior view.

**Figure S4.**
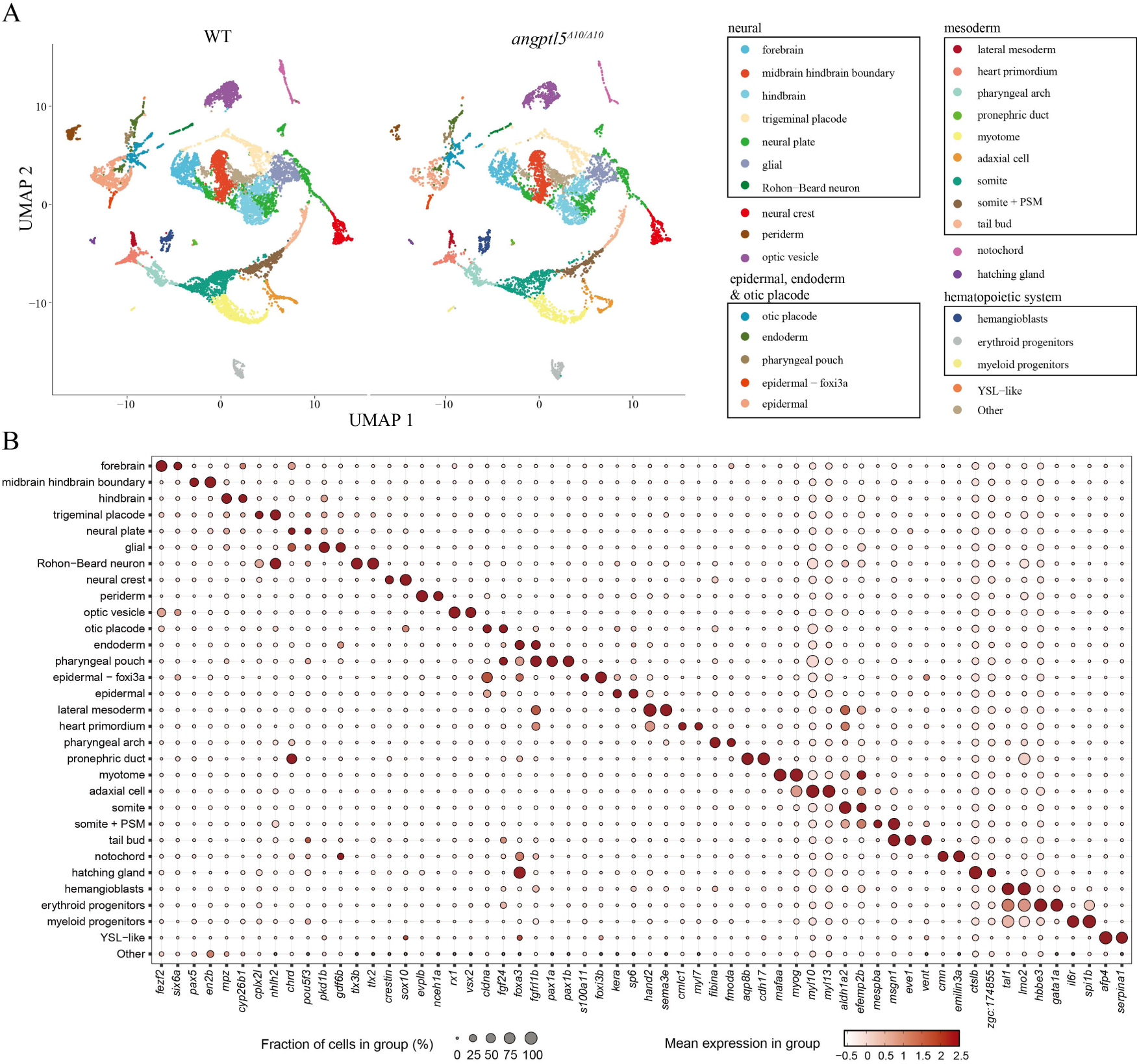
Integrative single cell RNA-seq analysis of 16 hpf WT and *angptl5^Δ10/Δ10^* embryos. (A) Integrative UMAP analysis of 16 hpf zebrafish embryos, with the UMAP results separated by the conditions of WT and *angptl5^Δ10/Δ10^*embryos. The WT dataset is from the published work (9). Each cell is coloured according to cell type annotations. (B) Dot plot showing the expression of two representative markers for each cluster. Colour represents the gene expression level, while dot size indicates the percentage of cells within the cluster expression the gene.

**Figure S5.**
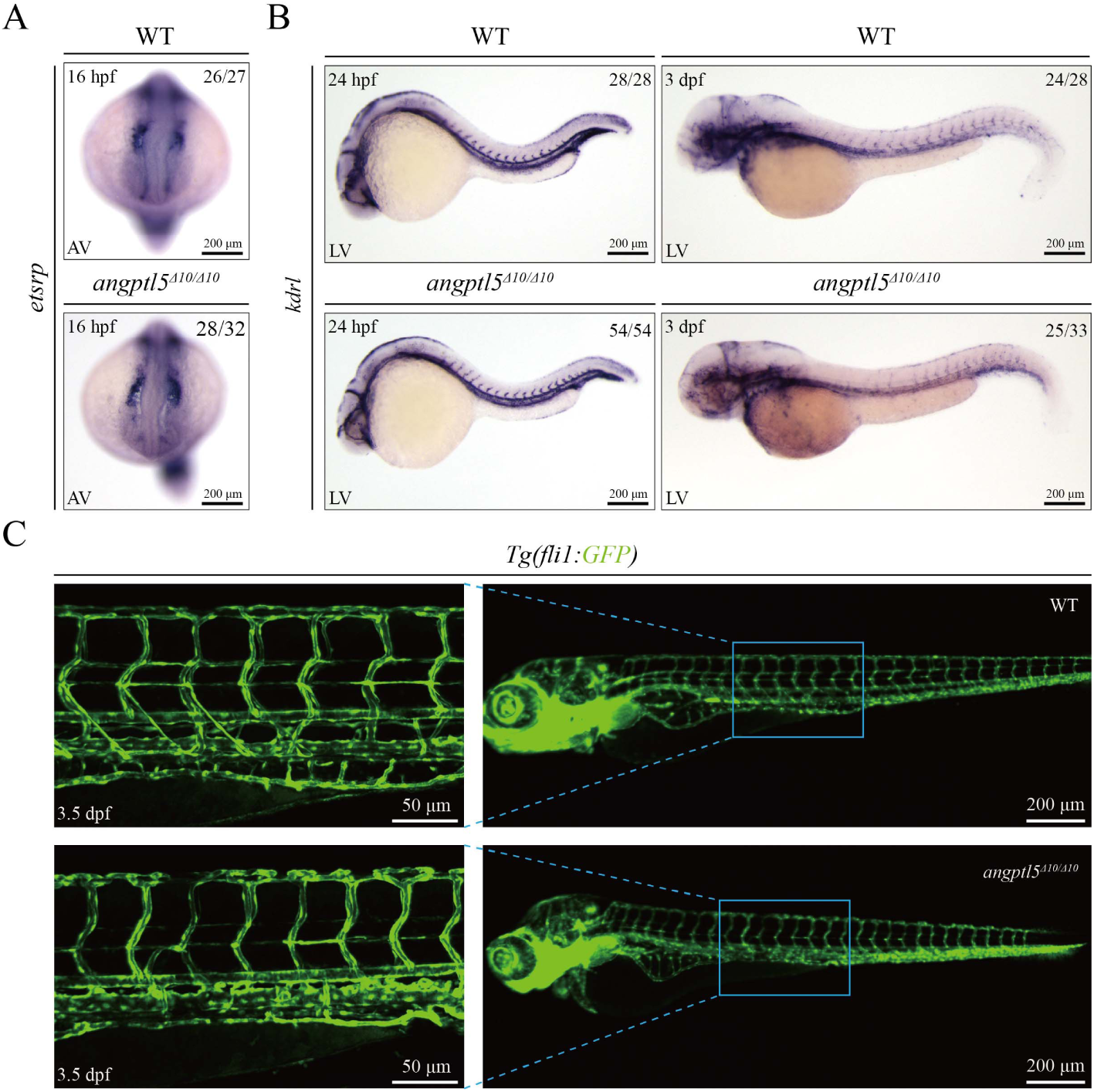
Comparison of primitive hematopoiesis and angiogenesis between WT and *angptl5^Δ10/Δ10^* embryos. (A-B) WISH of *etsrp* (A) and *kdrl* (B) in WT and *angptl5^Δ10/Δ10^* embryos. (C) Intersegmental blood vessels in WT and *angptl5^Δ10/Δ10^* embryos shown by *fli1:GFP* reporter. Each imaging was performed for at least three independent replicates. AV, anterior view; LV, lateral view.

**Figure S6.**
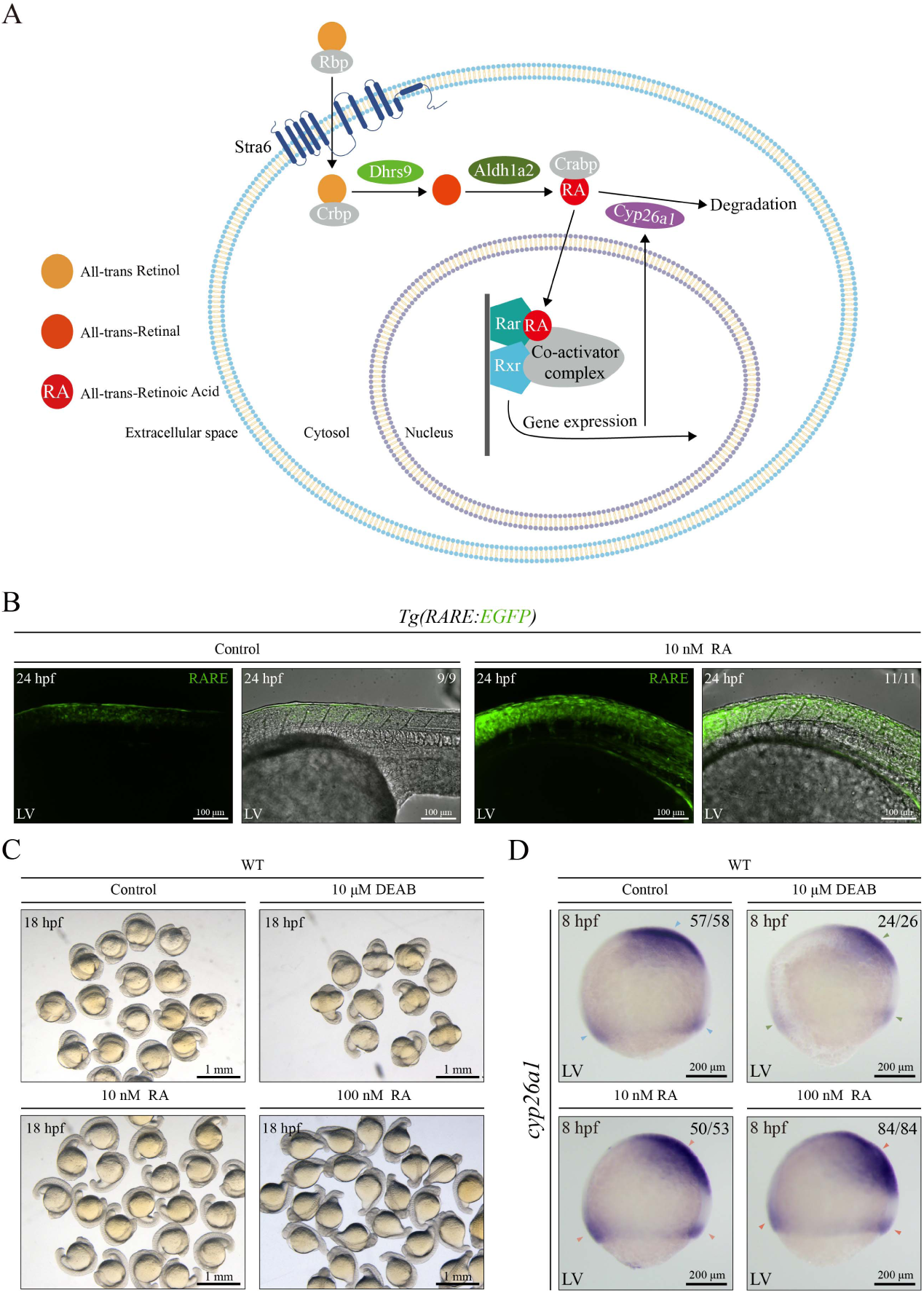
Retinoic acid signaling in early zebrafish development. (A) Schematic diagram of RA signaling pathway. (B) RA signaling in *Tg(RARE:EGFP)* embryos treated with RA from the shield stage to 24 hpf. Untreated embryos were used as control. (C) Morphology of WT embryos treated with RA inhibitor DEAB or different RA concentrations from the shield stage to 18-somite stage. Untreated embryos were used as control. (D) WISH of *cyp26a1* in WT embryos treated with DEAB or RA from the shield stage to 18-somite stage. Untreated embryos were used as control. LV, lateral view.

**Figure S7.**
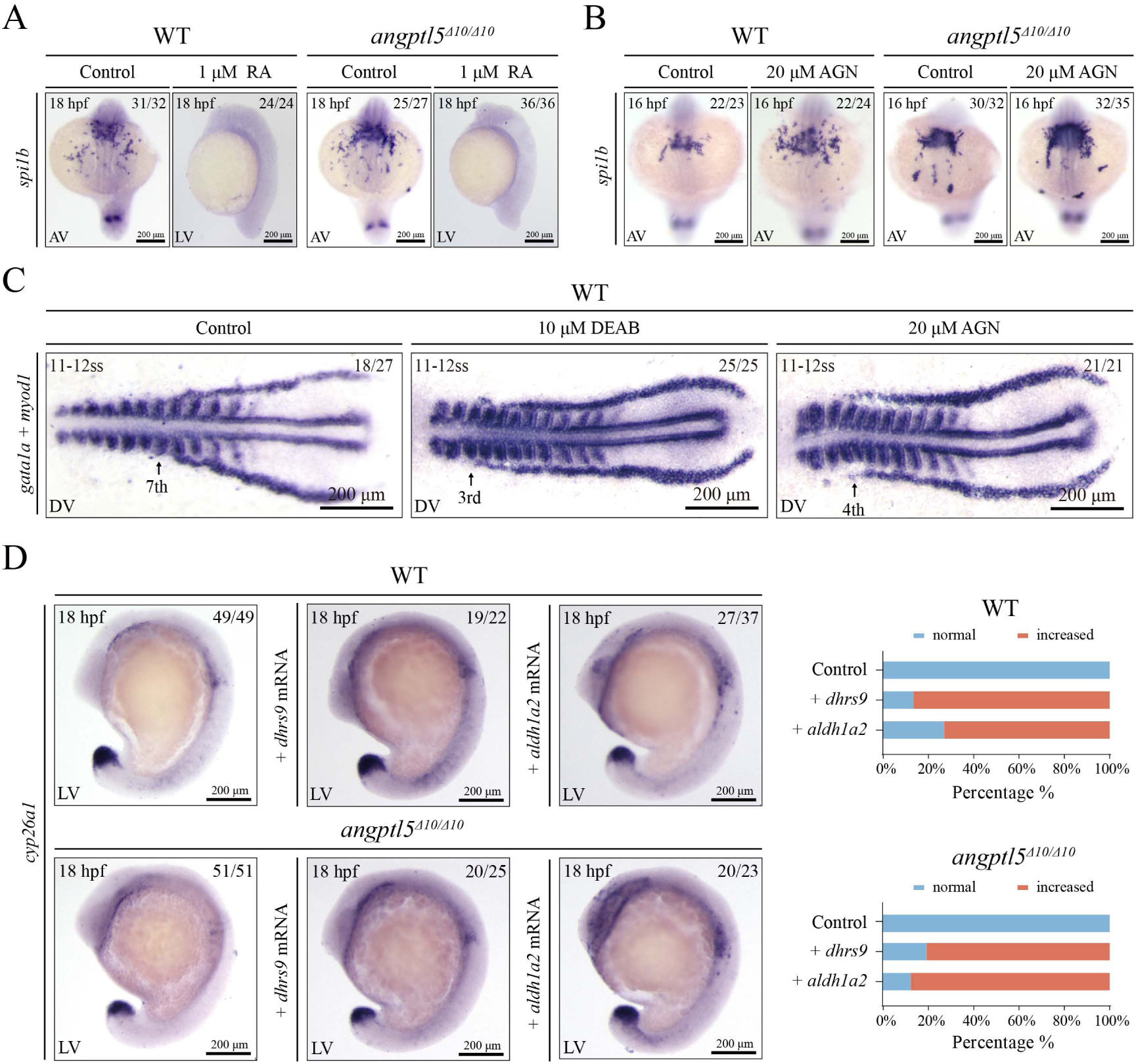
Regulation of *spi1b* and *gata1a* Expression by Retinoic Acid Signaling. (A-B) WISH of *spi1b* in WT and *angptl5^Δ10/Δ10^* embryos treated with RA (A) or RA inhibitor AGN (B). Untreated embryos were used as control. (C) WISH of *gata1a* and *myod1* in flat-mounted WT and *angptl5^Δ10/Δ10^* embryos treated with RA inhibitor DEAB or AGN. (D) WISH of *cyp26a1* in WT and *angptl5^Δ10/Δ10^*embryos injected with *dhrs9* or *aldh1a2* mRNA at the 1-cell stage. Uninjected embryos were used as control. Statistics are shown on the right of the representative photos. AV, anterior view; LV, lateral view.

**Figure S8.**
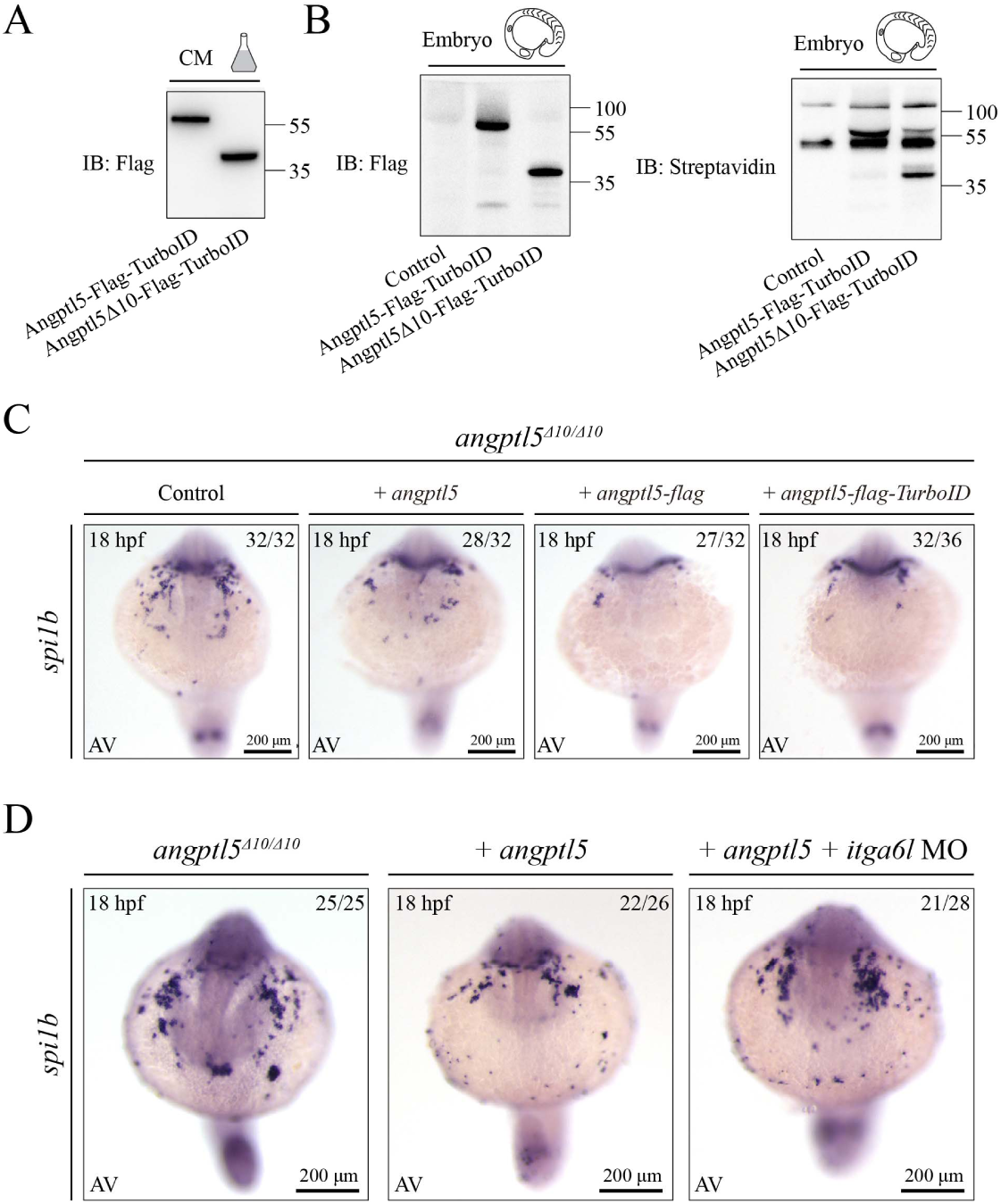
Secretion and functional analysis of tagged Angptl5. (A) Synthesis and secretion analysis of Angptl5-Flag-TurboID and Angptl5Δ10-Flag-TurboID in HEK293T cells after transfection with Angptl5-Flag-TurboID and Angptl5Δ10-Flag-TurboID plasmid for 48 hours. Conditioned medium (CM) were collected for western blot. (B) Expression of Angptl5-Flag-TurboID, Angptl5Δ10-Flag-TurboID and biotinylated proteins in zebrafish embryos. Embryos were injected with *Angptl5-Flag-TurboID* and *Angptl5Δ10-Flag-TurboID* mRNA at the 1-cell stage, and then embryos were collected for western blot at the 75% epiboly. (C) WISH of *spi1b* in *angptl5^Δ10/Δ10^*embryos injected with *angptl5*, *angptl5-flag* or *angptl5-TurboID* mRNA at the 1-cell stage. Uninjected embryos were used as control. (D) WISH of *spi1b* in *angptl5^Δ10/Δ10^* embryos. Embryos were injected with *angptl5* mRNA *± itga6l* MO at the 1-cell stage. Uninjected embryos were used as control. AV, anterior view.

**Figure S9.**
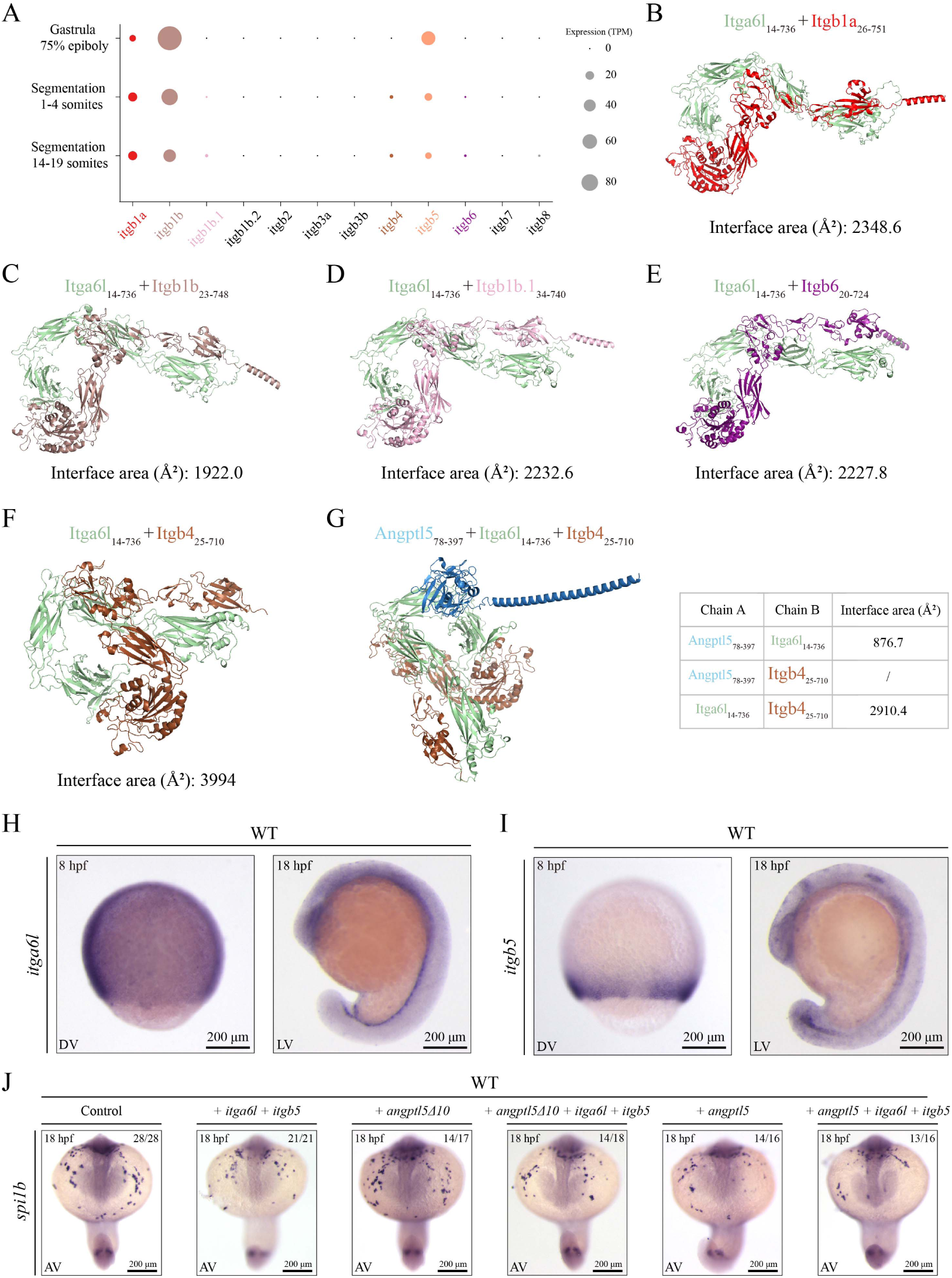
Interaction between Itga6l and integrin β-subunits. (A) Expression level of integrin β-subunits in zebrafish embryos from published work (13). (B-F) Structural models of the extracellular domain complex between Itga6l and: Itgb1a (B); Itgb1b (C); Itgb1b.1 (D); Itgb6 (E); Itgb4 (F). (G) Structural models of the Angptl5_78-397_-Itga6l_14-736_-Itgb4_25-710_ complexes. (H-I), WISH of *itga6* (H) and *itgb5* (I) in WT embryos. (J) WISH of *spi1b* in WT embryos injected with respective mRNA as indicated at the 1-cell stage. All structures were modelled using AlphaFold3. DV, dorsal view; LV, lateral view; AV, anterior view.

**Figure S10.**
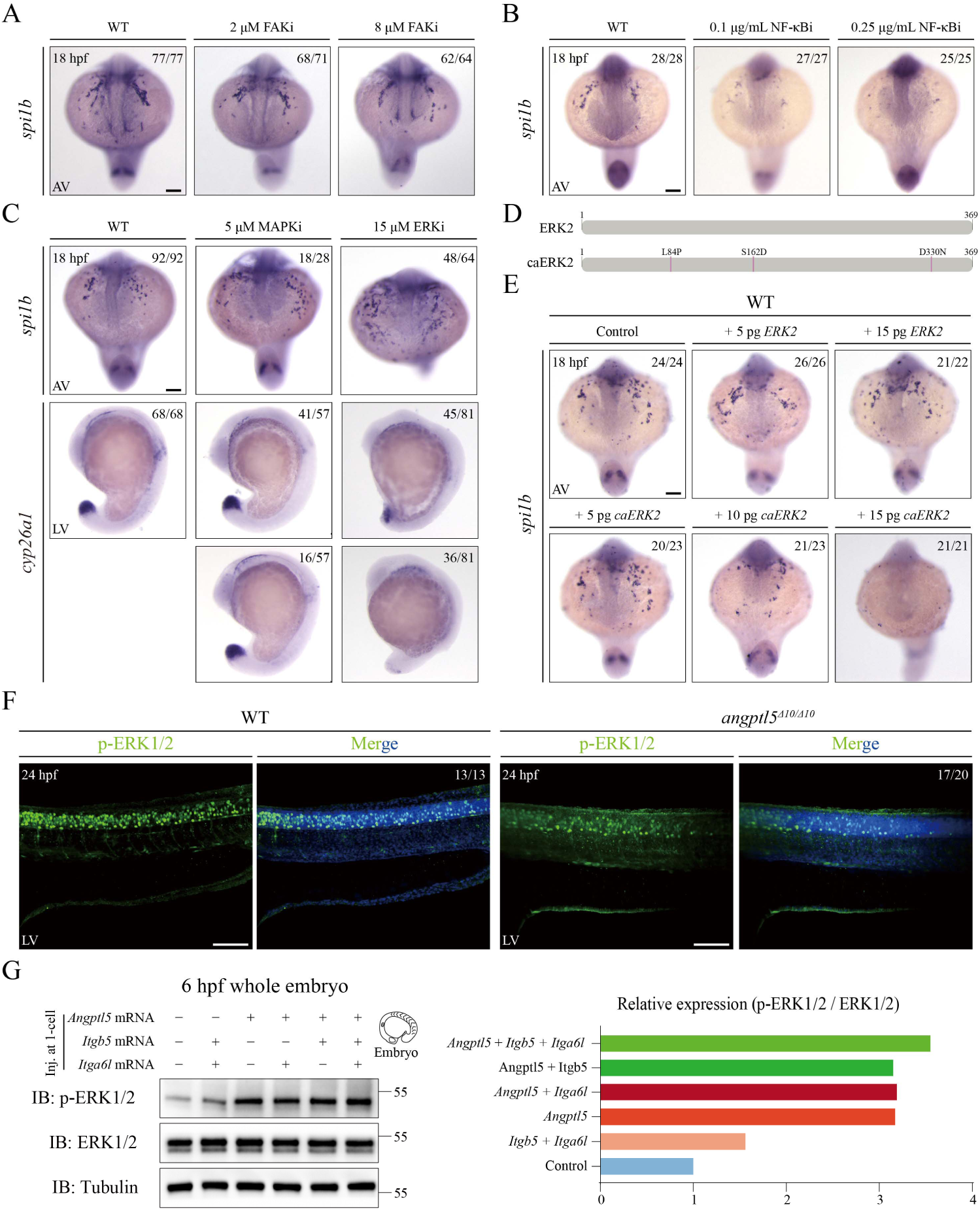
ERK signaling is reduced in *angptl5^Δ10/Δ10^* embryos. (A-C) WISH of *spi1b* in WT embryos treated with FAK inhibitor (A), NF-κB inhibitor (B), MAPK inhibitor and ERK inhibitor (C) from shield stage to 18-somite stage. Untreated embryos were used as control. (D) Schematic diagram of ERK2 and constitutively activated ERK2 (caERK2). (E) WISH of *spi1b* in WT embryos injected with *ERK2* or *caERK2* mRNA at the 1-cell stage. Uninjected embryos were used as control. (F) Projection of trunk region images from confocal stacks to show p-ERK1/2 in WT and *angptl5^Δ10/Δ10^* embryos by IF. (G) Expression of p-ERK in WT embryos injected with *angptl5* + *itga6l* + *itgb5* mRNA at the 1-cell stage. Statistics are shown on the right. AV, anterior view; LV, lateral view.

**Figure S11.**
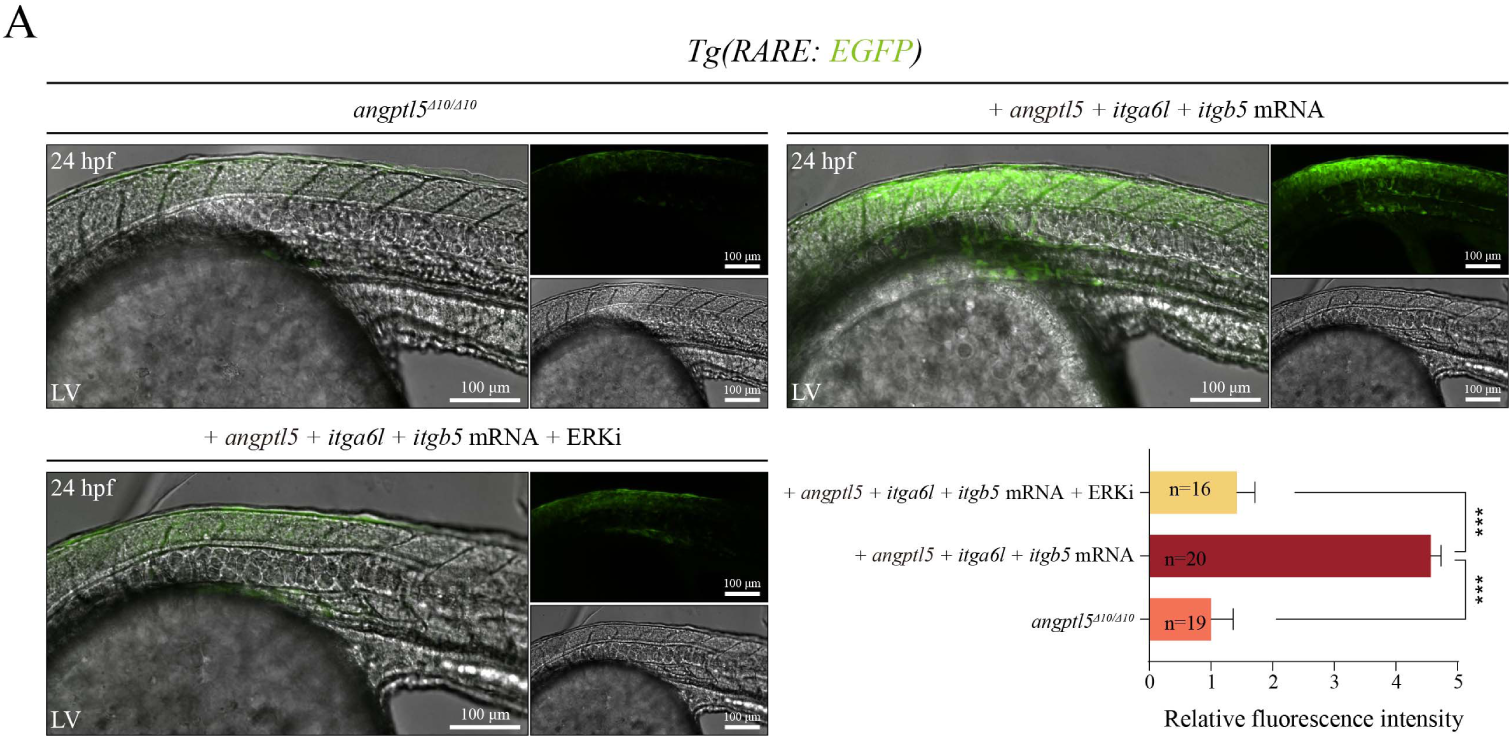
RA signaling is regulated by Angptl5-Integrin α6lβ5-ERK signaling cascade. (A) RA signaling in *angptl5^Δ10/Δ10^* embryos injected with *angptl5* + *itga6l* + *itgb5* mRNA at the 1-cell stage and treated with or without ERK inhibitor from the shield stage to 18-somite stage.

**Table S1.**
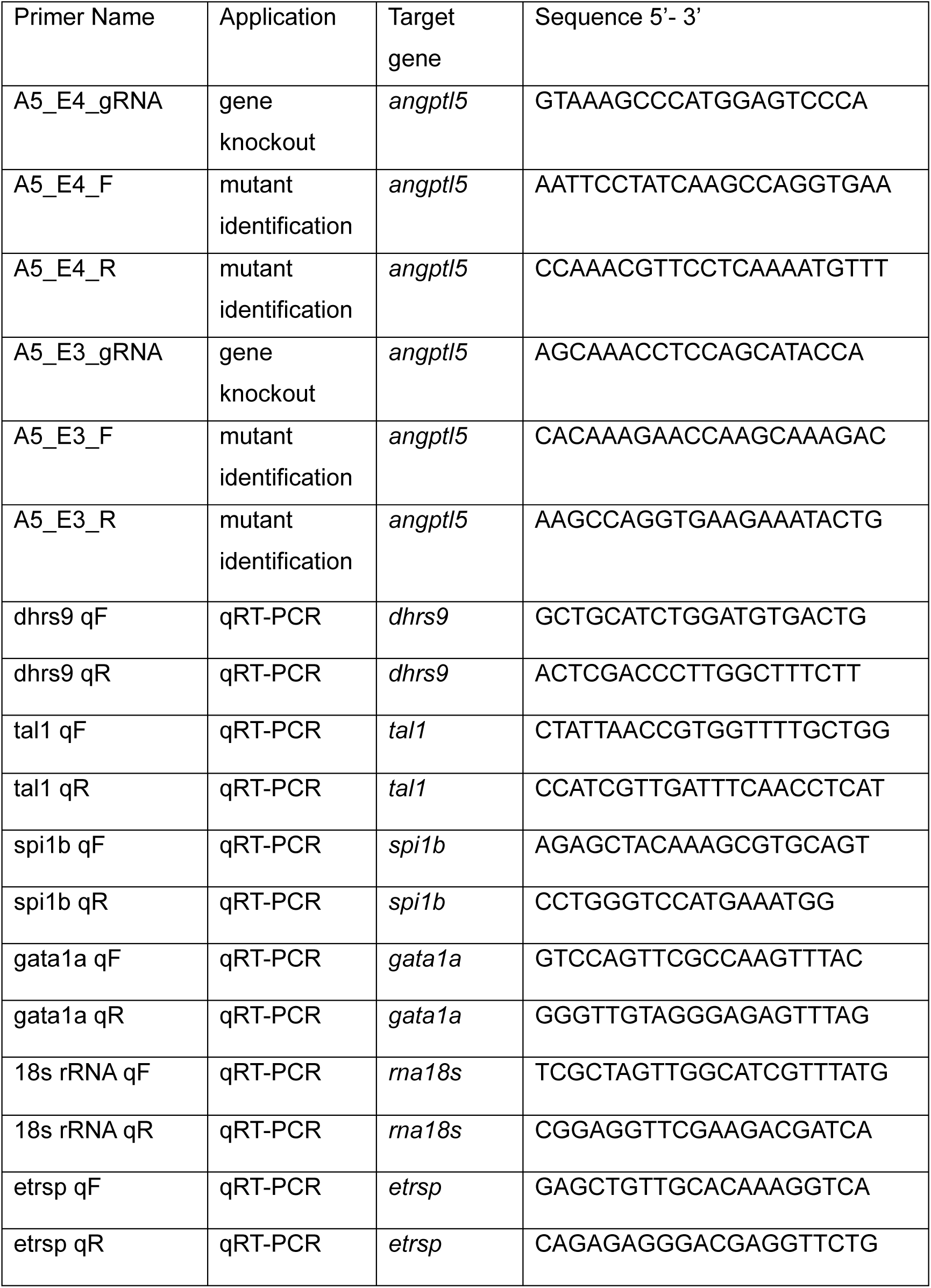
Primer sequences.

## References

1. H. W. Detrich, 3rd et al., Intraembryonic hematopoietic cell migration during vertebrate development. Proc Natl Acad Sci U S A 92, 10713–10717 (1995).

2. P. Herbomel, B. Thisse, C. Thisse, Ontogeny and behaviour of early macrophages in the zebrafish embryo. Development 126, 3735–3745 (1999).

3. C. M. Bennett et al., Myelopoiesis in the zebrafish, Danio rerio. Blood 98, 643–651 (2001).

4. L. J. Patterson et al., The transcription factors Scl and Lmo2 act together during development of the hemangioblast in zebrafish. Blood 109, 2389–2398 (2007).

5. K. Hsu et al., The pu. 1 promoter drives myeloid gene expression in zebrafish. Blood 104, 1291–1297 (2004).

6. J. L. Galloway, R. A. Wingert, C. Thisse, B. Thisse, L. I. Zon, Loss of gata1 but not gata2 converts erythropoiesis to myelopoiesis in zebrafish embryos. Developmental cell 8, 109–116 (2005).

7. J. Rhodes et al., Interplay of Pu.1 and gatal determines myelo-erythroid progenitor cell fate in zebrafish. Developmental Cell 8, 97–108 (2005).

8. R. Monteiro, C. Pouget, R. Patient, The gata1/pu.1 lineage fate paradigm varies between blood populations and is modulated by tif1γ. Embo J 30, 1093–1103 (2011).

9. R. K. T. Kam, Y. Deng, Y. L. Chen, H. Zhao, Retinoic acid synthesis and functions in early embryonic development. Cell and Bioscience 2 (2012).

10. J. L. de Jong et al., Interaction of retinoic acid and scl controls primitive blood development. Blood 116, 201–209 (2010).

11. M. R. Hawkins, R. A. Wingert, Zebrafish as a Model to Study Retinoic Acid Signaling in Development and Disease. Biomedicines 11 (2023).

12. M. E. Huang et al., Use of all-trans retinoic acid in the treatment of acute promyelocytic leukemia. Blood 72, 567–572 (1988).

13. J. Yang et al., Emerging roles of angiopoietin-like proteins in inflammation: Mechanisms and potential as pharmacological targets. J Cell Physiol 237, 98–117 (2022).

14. T. Kadomatsu, Y. Oike, Roles of angiopoietin-like proteins in regulation of stem cell activity. Journal of Biochemistry 165, 309–315 (2019).

15. Q. Yan et al., Interacts with Integrin α1β1 to Suppress HCC Angiogenesis and Metastasis by Inhibiting JAK2/STAT3 Signaling (Retracted article. See vol. 82, pg. 4299, 2022). Cancer research 77, 5831–5845 (2017).

16. S. Yumoto et al., Host ANGPTL2 establishes an immunosuppressive tumor microenvironment and resistance to immune checkpoint therapy. Cancer science 115, 3846–3858 (2024).

17. B. Aryal et al., ANGPTL4 deficiency in haematopoietic cells promotes monocyte expansion and atherosclerosis progression. Nat Commun 7 (2016).

18. C. C. Zhang, M. Kaba, S. Iizuka, H. Huynh, H. F. Lodish, Angiopoietin-like 5 and IGFBP2 stimulate ex vivo expansion of human cord blood hematopoietic stem cells as assayed by NOD/SCID transplantation. Blood 111, 3415–3423 (2008).

19. J. K. Zheng et al., Inhibitory receptors bind ANGPTLs and support blood stem cells and leukaemia development. Nature 485, 656–60 (2012).

20. D. E. Wagner et al., Single-cell mapping of gene expression landscapes and lineage in the zebrafish embryo. Science 360, 981–987 (2018).

21. A. Sur et al., Single-cell analysis of shared signatures and transcriptional diversity during zebrafish development. Developmental Cell 58 (2023).

22. C. Carbone et al., Angiopoietin-Like Proteins in Angiogenesis, Inflammation and Cancer. Int J Mol Sci 19 (2018).

23. T. Hato, M. Tabata, Y. Oike, The role of angiopoietin-like proteins in angiogenesis and metabolism. Trends Cardiovasc Med 18, 6–14 (2008).

24. B. Chestnut, S. Casie Chetty, A. L. Koenig, S. Sumanas, Single-cell transcriptomic analysis identifies the conversion of zebrafish Etv2-deficient vascular progenitors into skeletal muscle. Nat Commun 11 (2020).

25. H. Habeck et al., Analysis of a zebrafish VEGF receptor mutant reveals specific disruption of angiogenesis. Current Biology 12, 1405–1412 (2002).

26. N. Cabezas-Wallscheid et al., Vitamin A-Retinoic Acid Signaling Regulates Hematopoietic Stem Cell Dormancy. Cell 169, 807–823 e819 (2017).

27. C. O’Connor, P. Varshosaz, A. R. Moise, Mechanisms of Feedback Regulation of Vitamin A Metabolism. Nutrients 14 (2022).

28. Z. R. Xiong et al., In vivo proteomic mapping through GFP-directed proximity-dependent biotin labelling in zebrafish. Elife 10 (2021).

29. Y. A. Kadry, D. A. Calderwood, Chapter 22: Structural and signaling functions of integrins. Bba-Biomembranes 1862 (2020).

30. C. Kim, F. Ye, M. H. Ginsberg, Regulation of Integrin Activation. Annu Rev Cell Dev Bi 27, 321–45 (2011).

31. R. J. White et al., A high-resolution mRNA expression time course of embryonic development in zebrafish. Elife 6 (2017).

32. Y. Cui et al., CircHERC1 promotes non-small cell lung cancer cell progression by sequestering FOXO1 in the cytoplasm and regulating the miR-142-3p-HMGB1 axis. Mol Cancer 22, 179 (2023).

33. N. Mosca et al., LIM Homeobox-2 Suppresses Hallmarks of Adult and Pediatric Liver Cancers by Inactivating MAPK/ERK and Wnt/Beta-Catenin Pathways. Liver Cancer 11, 126–140 (2022).

34. J. H. Park et al., Asiatic acid attenuates methamphetamine-induced neuroinflammation and neurotoxicity through blocking of NF-kB/STAT3/ERK and mitochondria-mediated apoptosis pathway. J Neuroinflammation 14, 240 (2017).

35. H. Rian, S. G. Krens, H. P. Spaink, B. E. Snaar-Jagalska, Generation of Constitutive Active ERK Mutants as Tools for Cancer Research in Zebrafish. International Scholarly Research Notices 2013, 867613 (2013).

36. J. R. Brewer, P. Mazot, P. Soriano, Genetic insights into the mechanisms of Fgf signaling. Genes & Development 30, 751–771 (2016).

37. S. Romeo et al., Rare loss-of-function mutations in ANGPTL family members contribute to plasma triglyceride levels in humans. J Clin Invest 119, 70–79 (2009).

38. S. Kersten, Angiopoietin-like 3 in lipoprotein metabolism. Nat Rev Endocrinol 13, 731–739 (2017).

39. T. Kadomatsu, M. Endo, K. Miyata, Y. Oike, Diverse roles of ANGPTL2 in physiology and pathophysiology. Trends Endocrinol Metab 25, 245–254 (2014).

40. M. Endo, The Roles of ANGPTL Families in Cancer Progression. J UOEH 41, 317–325 (2019).

41. C. C. Zhang, M. Kaba, S. Iizuka, H. Huynh, H. F. Lodish, Angiopoietin-like 5 and IGFBP2 stimulate ex vivo expansion of human cord blood hematopoietic stem cells as assayed by NOD/SCID transplantation. Blood 111, 3415–3423 (2008).

42. M. Khoury et al., Mesenchymal Stem Cells Secreting Angiopoietin-Like-5 Support Efficient Expansion of Human Hematopoietic Stem Cells Without Compromising Their Repopulating Potential. Stem Cells Dev 20, 1371–1381 (2011).

43. J. Rhodes et al., Interplay of pu.1 and gata1 determines myelo-erythroid progenitor cell fate in zebrafish. Dev Cell 8, 97–108 (2005).

44. J. L. Galloway, R. A. Wingert, C. Thisse, B. Thisse, L. I. Zon, Loss of gata1 but not gata2 converts erythropoiesis to myelopoiesis in zebrafish embryos. Dev Cell 8, 109–116 (2005).

45. J. M. Frame, S. E. Lim, T. E. North, Hematopoietic stem cell development: Using the zebrafish to identify extrinsic and intrinsic mechanisms regulating hematopoiesis. Methods Cell Biol 138, 165–192 (2017).

46. A. Yen, M. S. Roberson, S. Varvayanis, A. T. Lee, Retinoic acid induced mitogen-activated protein (MAP)/extracellular signal-regulated kinase (ERK) kinase-dependent MAP kinase activation needed to elicit HL-60 cell differentiation and growth arrest. Cancer Res 58, 3163–3172 (1998).

47. S. D. Persaud, Y. W. Lin, C. Y. Wu, H. Kagechika, L. N. Wei, Cellular retinoic acid binding protein I mediates rapid non-canonical activation of ERK1/2 by all-trans retinoic acid. Cell Signal 25, 19–25 (2013).

48. F. Zassadowski et al., Lithium chloride antileukemic activity in acute promyelocytic leukemia is GSK-3 and MEK/ERK dependent. Leukemia 29, 2277–2284 (2015).

## SI References

1. C. Lawrence, The husbandry of zebrafish (Danio rerio): A review. Aquaculture 269, 1–20 (2007).

2. C. B. Kimmel, W. W. Ballard, S. R. Kimmel, B. Ullmann, T. F. Schilling, Stages of embryonic development of the zebrafish. Developmental Dynamics 203, 253–310 (1995).

3. B. Thisse, C. Thisse, In situ hybridization on whole-mount zebrafish embryos and young larvae. Methods Mol Biol 1211, 53–67 (2014).

4. A. Urasaki, G. Morvan, K. Kawakami, Functional dissection of the Tol2 transposable element identified the minimal cis-sequence and a highly repetitive sequence in the subterminal region essential for transposition. Genetics 174, 639–649 (2006).

5. T. Cheng, Y. Y. Xing, Y. Dong, P. F. Xu, Protocol for generation and assessment of head-like structure in zebrafish. STAR Protoc 4, 102553 (2023).

6. N. N. Chang et al., Genome editing with RNA-guided Cas9 nuclease in Zebrafish embryos. Cell Res 23, 465–472 (2013).

7. A. L. van Boxtel et al., A Temporal Window for Signal Activation Dictates the Dimensions of a Nodal Signaling Domain. Developmental Cell 35, 175–185 (2015).

8. Y. H. Hao et al., Dictionary learning for integrative, multimodal and scalable single-cell analysis. Nat Biotechnol 42 (2024).

9. A. Sur et al., Single-cell analysis of shared signatures and transcriptional diversity during zebrafish development. Developmental Cell 58 (2023).

10. X. X. Fu et al., A spatiotemporal barrier formed by Follistatin is required for left-right patterning. Proceedings of the National Academy of Sciences of the United States of America 120, e2219649120 (2023).

11. Z. R. Xiong et al., In vivo proteomic mapping through GFP-directed proximity-dependent biotin labelling in zebrafish. Elife 10 (2021).

12. L. Yan et al., Maternal Huluwa dictates the embryonic body axis through beta-catenin in vertebrates. Science 362 (2018).

13. R. J. White et al., A high-resolution mRNA expression time course of embryonic development in zebrafish. Elife 6 (2017).

